# Coarse-grained models for simulations of double-stranded nucleic acids for mixed protein–nucleic acid condensates

**DOI:** 10.64898/2026.08.14.744942

**Authors:** Ikki Yasuda, Giulio Tesei, Eiji Yamamoto, Kenji Yasuoka, Kresten Lindorff-Larsen

## Abstract

Biomolecular condensates function as membraneless compartments, and some protein condensates can selectively concentrate single-stranded nucleic acids while excluding double-stranded nucleic acids. Understanding how nucleic acid structure affects partitioning into condensates has important implications for nucleic acid activity and function within condensates. Here, we present a set of coarse-grained two-bead-per-nucleotide models for simulations of double-stranded RNA and DNA in the CALVADOS framework. Our models separately represent the backbone and base, and maintain the helical structures using an elastic network potential tuned to capture chain stiffness. For dsRNA, the base stickiness was tuned using experimental data on differential partitioning of single- and double-stranded RNA into Ddx4N1 condensates in order to account for reduced base accessibility upon duplex formation. This RNA structural selectivity varied with the balance of electrostatic and non-electrostatic interactions, as revealed by simulations of condensates of the CAPRIN1 disordered region at varying ionic concentrations and with an R-to-K sequence variant. Finally, we developed parameters for double-stranded DNA using a similar approach. We envision that the CALVADOS models for double-stranded RNA and DNA will be useful for studying co-condensates of proteins and structured nucleic acids.

## 1 Introduction

Cellular phase separation is a key mechanism of compartmentalisation, generating biomolecular condensates that regulate diverse cellular functions and contribute to disease development.^1–4^ These condensates create unique physical and chemical environments distinct from the surrounding nucleoplasm and cytoplasm. Condensates not only concentrate biomolecules but are also associated with their conformational stability, such as changes in the stability and conformational equilibria of globular proteins within condensates observed in experiments^5,6^ and computation,^7,8^ conformational-change-induced condensation,^9,10^ and destabilisation of hybridised and aggregated nucleic acids.^11–13^ Such structural rearrangements can be central to the function of condensates, modulating catalytic activity and signalling of biomolecules.^14^ While thermodynamic linkage relationships provide a theoretical framework for quantifying the coupling between changes in solvent environment and conformational changes,^15,16^ we lack a quantitative framework for understanding these structural effects at the molecular level.^17^

The inner environment of condensates can modulate the conformational stability of nucleic acids within protein–RNA mixed condensates.^18,19^ In a seminal study, Nott *et al.* showed that condensates of the intrinsically disordered region (IDR) of DEAD-box helicase 4 (Ddx4), a key component of P granules, recruit single-stranded (ss) nucleic acids substantially more strongly than double-stranded (ds) forms.^11^ The difference in partitioning free energies between ds- and ss-nucleic acids into the condensates can be connected to the difference in hybridisation free energies inside and outside the condensate environment through a thermodynamic cycle, showing that the ds-form is destabilised relative to the ss-form in Ddx4N1 condensates.^12^ Similarly, the enhanced dissociation of dsRNA has been reported in condensates of the C-terminal IDR of CAPRIN1.^20^ Choi *et al.* used short-peptide model systems of multiphasic phase separation comprising polyR/K/D, showing that ssRNA is more enriched in the arginine-rich inner layers than dsRNA, indicating that the arginine-rich inner layer stabilised ssRNA more than the lysine-rich outer layer.^21^ Complementing these experiments, Boccalini *et al.* performed atomistic simulations of short RNAs within peptide condensates and demonstrated that base atoms in unfolded conformations form hydrogen bonds with the surrounding peptides, whereas such interactions are reduced when intramolecular base pairing occurs.^22^ These experimental and computational studies suggest that base accessibility plays a key role in the differential partitioning of ss- and ds-nucleic acids into protein condensates.

This selective nucleic acid recruitment into condensates can be driven by both site-specific interactions and chemistry-specific interactions. Site-specific interactions form on complementary surfaces, yielding strong binding affinity.^23^ These interactions, including RNA interactions with RNA-binding proteins, can drive protein condensation with specific RNAs.^24–26^ In contrast, chemistry-specific interactions arise from complementary chemical groups, and depend less precisely on local structure.^23^ For nucleotides, these interactions involve the negatively charged phosphate backbone and the *π* interactions of bases. ^27–29^ Partitioning of ssRNA is not highly specific to pairs of IDRs and nucleic acids, ^12,20,21^ and therefore the differences in chemistry-specific interactions between the ss/ds forms likely govern their behaviour in condensate systems.

The molecular structure and chemistry of nucleic acids influence thermodynamic properties within condensates, such as protein–nucleic acid interactions, disruption of protein interaction networks upon insertion of nucleic acids, and the conformational free energy landscape of nucleic acids.^12,30^ These factors are ideally integrated in molecular simulations in which both condensation and conformational changes are captured. However, atomistic simulations are often limited by the available timescales and system sizes for condensate systems.^22,31^ Alternatively, coarse-grained models that capture both the nucleic acid structure and the sequence-dependent chemistry of amino acids and nucleotides could address conformation-dependent partitioning into condensates.

Coarse-grained molecular models have been developed and used for various nucleic acid systems, such as tetraloops, hairpins, flexible single-stranded and helical duplex nucleic acids, ribozymes, and riboswitches.^32–34^ These models have been designed to reproduce properties related to structure,^35–38^ thermodynamics,^39,40^ mechanical responses,^41–43^ and intra- and intermolecular interactions of protein–RNA complexes. ^44–46^ Moreover, RNAs can undergo condensation and aggregation without proteins.^47^ Nguyen *et al.* developed a single-site-per-nucleotide model that treated Watson-Crick base-pairing interactions for low-complexity RNA repeats to predict their folding–unfolding transition and condensation.^48^ Similarly, Tejedor *et al.* developed a two-bead-per-nucleotide model that well captured the sequence dependence of the hybridisation and condensation.^49^ Similar single-site-per-nucleotide models were also developed to represent DNA structures in nucleosomes by integrating atomistic simulation and experimental data,^50^ and to predict the thermodynamic stability of RNA under mechanical tension.^51^

In parallel, we and others have developed coarse-grained models at the nucleotide resolution for simulation of phase separation with proteins,^52–56^ complementing super-coarsegrained models^57,58^ and intermediate-resolution models.^59–61^ The nucleotide-resolution models use explicit charges and chains and often assume unstructured nucleotides, though some models have been designed for simulation of structured nucleic acids.^62–64^ In simulations of proteins, models developed for disordered proteins have been shown to require adaptation for application to folded domains, either by changing the energy scale depending on how buried residues are^65–67^ or by shifting the coarse-grained representation geometry,^7^ suggesting the necessity of similar adaptations for dsRNA models. ^68^ Therefore, there is a remaining challenge in integrating RNA structural models with protein–RNA phase-separation models in a single framework.

Here, we present models of dsRNA and dsDNA in the CALVADOS framework for simulations of mixed protein–nucleic acid condensates. Our dsRNA model is built upon the CALVADOS framework,^7,56,69–71^ with the protein model related to other one-bead-per-residue models.^54,55,72–75^ Such models describe interactions between macromolecules using electrostatics and non-electrostatic interactions (also referred to as stickiness), with solvents and ions treated implicitly. We previously developed CALVADOS models for disordered proteins,^69,70^ proteins with both ordered and disordered regions,^7,71^ and a two-bead-per-nucleotide model for flexible sequence-independent ssRNA.^56^

One potential advantage of a two-bead-per-nucleotide representation is that it can separate electrostatic and stickiness interactions from the backbone and bases, which we suggest is also useful for representing surface chemistry of helical nucleic acids. Nevertheless, because these simple coarse-grained models do not maintain tertiary structures unless constrained using an additional potential, the CALVADOS models for double-stranded RNA and DNA employ a structure-based constraint and recalibrated interaction parameters. Here, we focused our efforts on defining elastic network potentials to maintain the helical structures and on the stickiness parameter of the base bead. We calibrated the interaction parameter of the base against the experimental measurements of differential partitioning free energies of ssRNA and dsRNA into Ddx4N1 condensates,^12^ using a tie-line interpolation approach that enables more direct comparison of simulation and experimental conditions. The force constant for the elastic network force was parametrised against the persistence length of dsRNA. Furthermore, we demonstrate how the balance between electrostatic and non-electrostatic interactions affects ssRNA and dsRNA partitioning using CAPRIN1 condensate systems. Following the development of this dsRNA model, we applied a similar approach to develop a dsDNA model.

## 2 Results

### 2.1 Overview of the CALVADOS model for double-stranded RNA

We developed a dsRNA model, which represents one nucleotide by two beads, corresponding to the backbone and base (Figs. 1a and b), consistent with our previously developed CALVADOS–RNA model for flexible ssRNA.^56^ Our dsRNA model describes dsRNA in an A-form double-stranded structure (Fig. 1a). Because of the limited chemical specificity at this coarse-grained resolution and the scarcity of experimental data for model parametrisation,^28,29^ our models are simple sequence-independent representations that capture electrostatics at the backbone and stickiness of bases. Although terminal nucleotides in dsRNA are more flexible^76^ and potentially more interactive than internal nucleotides, this difference is not considered in our current model.

**Figure 1:**
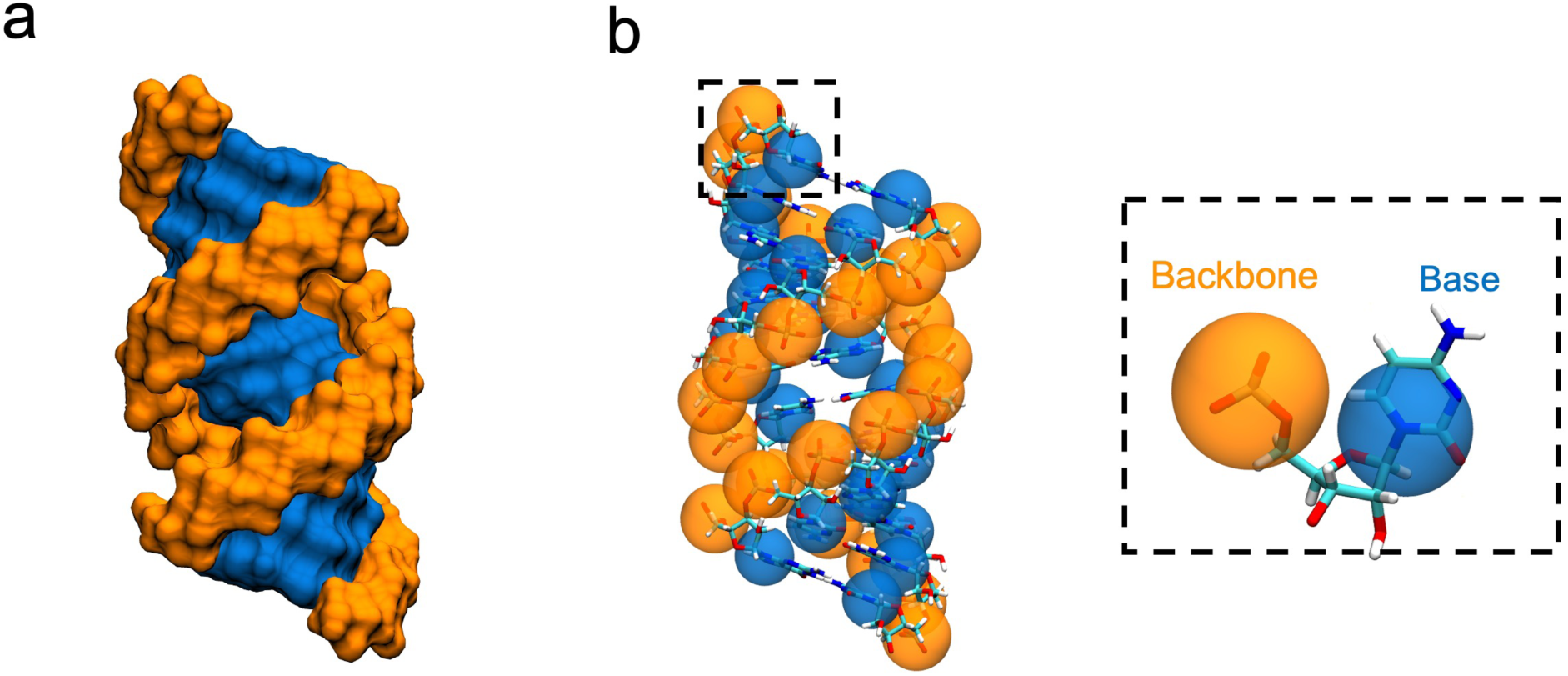
Representation of double-stranded RNA (dsRNA) using CALVADOS-dsRNA. (a) Atomistic structure of A-form dsRNA shown using a surface representation. Backbone and base atoms are coloured in orange and blue, respectively. (b) Coarse-grained representation using CALVADOS-dsRNA. The atomistic structure is overlaid in a stick representation. Backbone and base beads are coloured in orange and blue, respectively.

Our dsRNA model contains bonded terms describing bonds, angles, and local stacking interactions, and non-bonded terms for electrostatic interactions and stickiness (described via the Ashbaugh-Hatch (AH) potential). In addition, the model includes an elastic network to maintain the folded structure of dsRNA (see also Methods). The differences from CALVADOS-RNA are that: (1) the AH stickiness parameter, *λ*, of the base bead is reduced, and (2) an elastic network is introduced to maintain the folded structures, as used in other higher-resolution models,^59,60^ and the CALVADOS 3 model for proteins.^7^

Because the accessibility of bases differs between ssRNA and dsRNA, we assign different parameters to the base beads in the two forms. A similar strategy of using structure-dependent parameters has been adopted in other coarse-grained protein models.^54,65,73,77^ The bases in ssRNA can participate in hydrogen bonding, hydrophobic contacts, *π*-stacking, and *π*–cation interactions in protein–RNA interactions. ^28,78^ We previously assigned a high stickiness parameter to the base beads in our ssRNA model (*λ* = 1.18) from our calibration against the SAXS-derived radius of gyration of flexible homotypic RNAs.^56^ In contrast, in dsRNA, interaction sites of the bases are partly occupied by the complementary strand.^21,22^ In our dsRNA representation, the separation of the backbone and base beads plays a role similar to the centre-of-mass representation that we introduced for proteins to improve the description of the molecular surface.^7^ However, our dsRNA representation still leaves the base beads more exposed than the corresponding atomistic bases, and the exposed atom groups are predominantly polar. Accordingly, we used a lower *λ* value for the base beads in the dsRNA model.

The conformational flexibility in coarse-grained models can be modulated by the strength of the elastic network,^79^ and thus we tuned the force constant for this term to represent the stiffness of dsRNA. Since a relatively short (24 base pairs, bp) RNA was used in the partitioning experiment, we first modelled dsRNA as a rigid body to tune the base stickiness. Subsequently, we tuned the elastic force constant using longer dsRNA chains, and validated the results using the partitioning behaviour.

### 2.2 Partitioning of ssRNA into Ddx4N1 condensates

As a reference for our dsRNA model, we first investigated the partitioning of ssRNA into condensates of the N-terminal IDR of Ddx4 (hereafter Ddx4N1) modelled by combining CALVADOS 2 and CALVADOS-RNA. Ddx4N1 undergoes condensation at physiological salt concentrations both *in vitro* and *in vivo*,^80^ and selectively recruits ssRNA.^11,12^ We modelled the 24-nucleotide (nt) CAU repeat sequence used by Nott *et al.*,^12^ treating it as a disordered chain. We performed coarse-grained molecular simulations of Ddx4N1 and ssRNA, demonstrating ssRNA partitioning into Ddx4N1 condensates (Figs. 2a and b). Because the experimental bulk concentrations of both Ddx4N1 and ssRNA are too low to be conveniently accessible in simulations, we varied the relative amounts of Ddx4N1 and ssRNA to construct a phase diagram.

**Figure 2:**
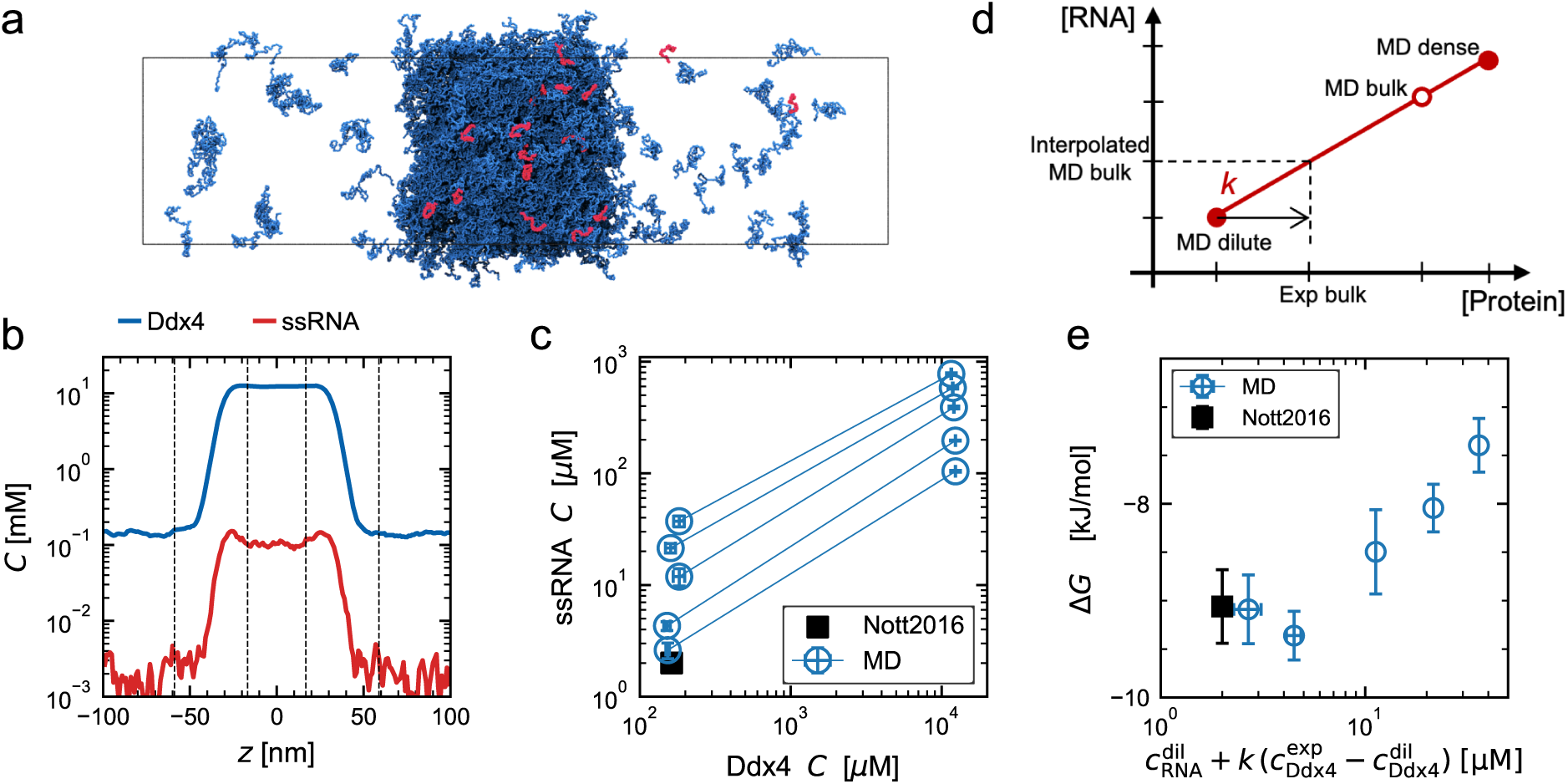
Partitioning of 24-nt single-stranded RNA (ssRNA) into Ddx4N1 (Ddx4) condensates. (a) A snapshot of the simulation system containing Ddx4N1 and 24-nt ssRNA. For clarity, ssRNA is shown in the front periodic image, while Ddx4N1 is shown in the unit cell. (b) Density profiles of Ddx4N1 and ssRNA with black dashed lines indicating the boundaries of the dense-phase, interfacial, and dilute-phase regions. (c) Phase diagram of the mixture of Ddx4N1 and ssRNA. The concentrations used in the experiments by Nott *et al.*^12^ are shown as a black square. Error bars represent the standard error of the mean estimated using the time-blocking method or the standard error of the mean over multiple replicas. (d) Illustration of tie-line interpolation to obtain an interpolated RNA concentration at an experimental protein concentration. *k* denotes the slope of the tie-line. (e) Partitioning free energy, Δ*G*, with varying ssRNA concentrations at a fixed Ddx4N1 concentration of 160 *µ*M. RNA concentration on the *x* axis is estimated using tie-line interpolation.

Concentrations of the dilute and dense phases were computed from the density profiles (Figs. 2b and S1), and used to construct the phase diagram of Ddx4N1 and ssRNA. The positive slope of the tie-lines indicates associative condensation of Ddx4N1 and ssRNA (Fig. 2c).^81^ Over the range of concentrations of ssRNA that we studied, the Ddx4N1 concentrations remained almost constant in both phases, indicating favourable condensation of Ddx4N1 via homotypic interactions.

We then used the phase diagram to match RNA concentrations between experimental and simulation conditions. For simplicity, we approximated the experimental system as coexisting dense and dilute phases, which is reasonable for micron-sized droplets where the volumes of interface regions are negligible. Under this two-phase approximation, when the experimental bulk concentrations (160 *µ*M Ddx4N1 and 2 *µ*M ssRNA) lie on the tie-line obtained from our simulations, the corresponding predicted dilute- and dense-phase concentrations can be compared to those from the experimental condition.

We introduced a tie-line interpolation to estimate the ssRNA concentration at the experimental Ddx4N1 concentration (Fig. 2d),

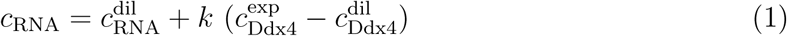

where 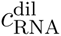 and 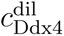 are the dilute-phase concentrations of RNA and Ddx4N1 in simulations, respectively. 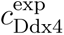 is the experimental Ddx4N1 concentration. *k* is the tie-line slope,^81–84^ defined as 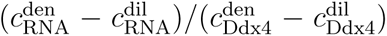, where 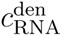 and 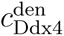 are the dense-phase concentrations of RNA and Ddx4N1. Because the surface volume in MD systems cannot be ignored, the bulk concentrations may deviate from the tie-line. Therefore, the dilute phase concentration was used as the reference in Eq. 1. This interpolation enables more direct comparison of RNA concentrations in simulations and experiments.

We applied this tie-line interpolation to compare our simulations to the experiments. In practice, the correction terms were small because the dilute-phase concentration of Ddx4N1 in our simulations was close to the experimental value (Fig. 2c). In cases where 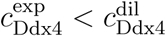 we attributed the discrepancy to uncertainty in sampling of dilute phase concentrations and extrapolated along the tie-line accordingly.

Next, we quantified the ssRNA partitioning into Ddx4N1 condensates using the partitioning free energy, defined as

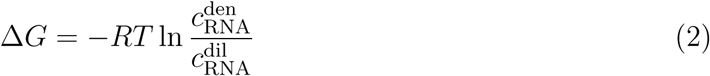

with the gas constant *R* and temperature *T*, and investigated the effect of ssRNA concentration. In our simulations, we found an increase in Δ*G* that was proportional to the ssRNA concentration (Fig. 2e), which we attribute to a crowding effect of ssRNA within the condensates. The lowest RNA ratio system yielded an interpolated ssRNA concentration of 2.6 ± 0.4 *µ*M and Δ*G*_ss_ of −9.1 ± 0.4 kJ*/*mol, which agrees with the experiment within uncertainty (Tab. S1). These results indicate that our ssRNA model captures ssRNA partitioning into Ddx4N1 condensates with reasonable accuracy.

### 2.3 dsRNA exclusion requires reduced base stickiness

To examine how differences between ssRNA and dsRNA affect their partitioning behaviour, we performed simulations of Ddx4N1 condensates with dsRNA (Fig. 3a). Our dsRNA model assumes that the dsRNA duplex remains intact within the condensate. This assumption is supported by the predicted stability of the 24-bp dsRNA duplex within condensates, whose stabilisation free energy in solution (∼ −203 kJ*/*mol) remains sufficiently large for the duplex to stay almost fully hybridised even after accounting for condensate-induced destabilisation.^12^ Consistent with this view, Ueno *et al.* reported that annealed 20-bp oligonucleotides remained hybridised within Ddx4N1 droplets on a timescale of approximately 1 hour.^85^ Although the FRET measurements by Nott *et al.* suggested structural variations of the duplex within condensates,^12^ treating dsRNA as a stable duplex in our simulations is expected to be a reasonable approximation.

**Figure 3:**
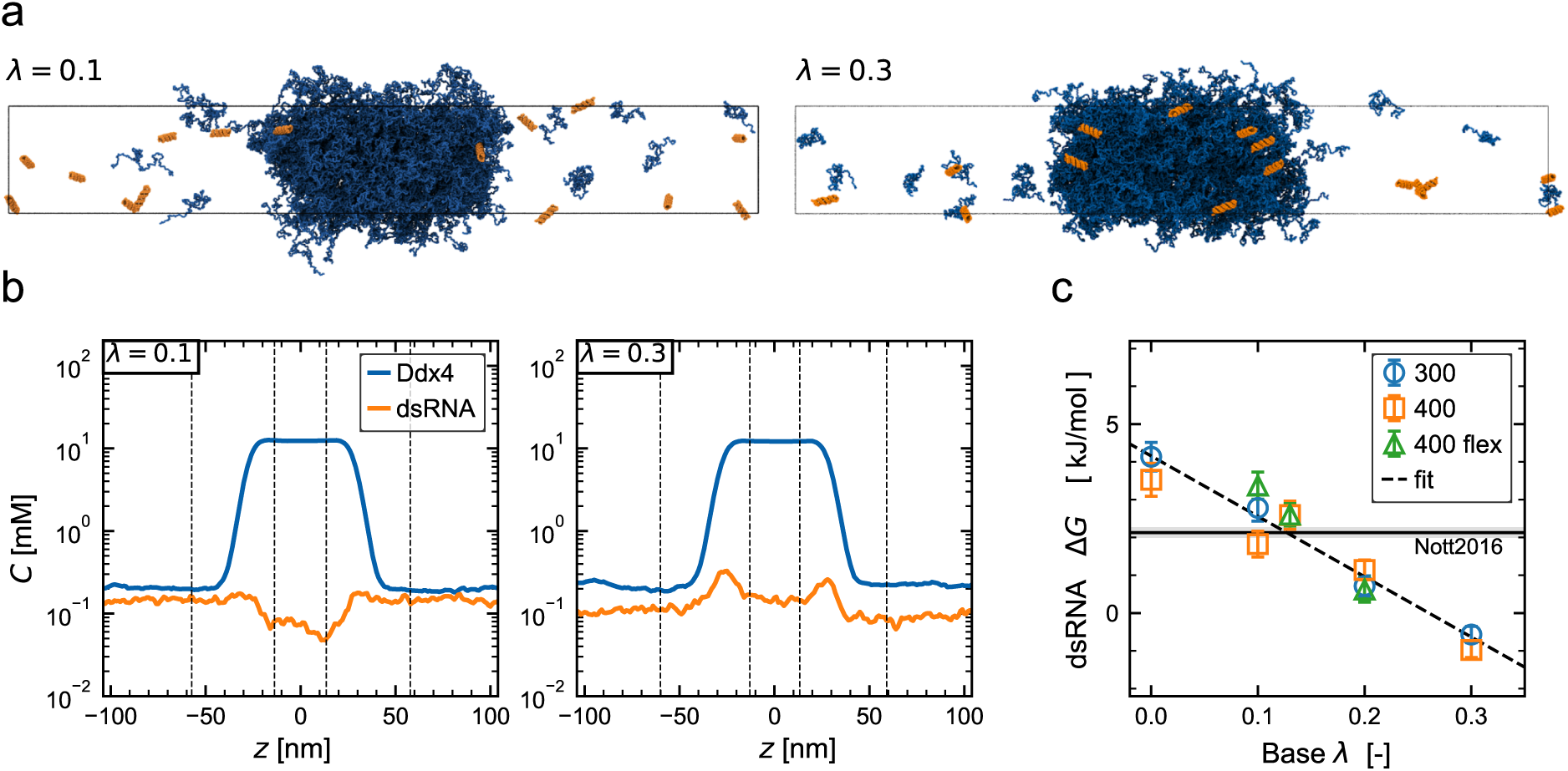
Partitioning of 24-bp dsRNA into Ddx4N1 condensates. (a) Snapshots of systems using base stickiness parameters, *λ* = 0.1 and 0.3. Ddx4N1 and dsRNA are coloured in blue and orange, respectively. For clarity, dsRNA is shown in the front periodic image, while Ddx4N1 is shown in the unit cell. (b) Density profiles corresponding to (a). Black dashed lines indicate boundaries of the dense-phase, interfacial, and dilute-phase regions. (c) Relationship between the base *λ* and partitioning free energy of dsRNA, Δ*G*. Labels indicate simulation conditions: ‘300’ and ‘400’ denote the number of Ddx4N1 chains in the simulation boxes using a rigid dsRNA model; ‘400 flex’ denotes 400 Ddx4N1 chains using a flexibility-tuned dsRNA model. The black dashed line shows a linear regression model fit to all data points with resulting slope −16.0 kJ*/*mol and intercept 4.2 kJ*/*mol. The black solid line with shaded region represents the experimental Δ*G*.

We constrained the dsRNA structure with a strong elastic network potential, and varied the *λ* (stickiness) value of the base of dsRNA from 0 to 0.3 to weakly exclude dsRNA (for comparison, the base parameter of ssRNA is 1.18). In these simulations we used fixed Ddx4N1 and dsRNA concentrations. In the Ddx4N1–ssRNA systems, the ssRNA concentration affected the partitioning free energy, an effect we attribute to ssRNA accumulation in the condensates. In contrast, when using base *λ* values in the range of 0–0.3 for dsRNA, we observed very little localisation of dsRNA into the condensates (Figs. 3b and S2; dsRNA concentration ∼ 0.1 mM). Therefore, we assumed that the effect of dsRNA accumulation was negligible (see validation below). The thickness of the condensates was sufficient to obtain the bulk region of the dense phase. Additionally, we observed surface accumulation of dsRNA at a moderate stickiness (*λ* = 0.3), but not at *λ* = 0.1.

Subsequently, we obtained the partitioning free energy of the dsRNA into the Ddx4N1 condensates. The relationship between the base *λ* and Δ*G* shows an approximately linear decrease in Δ*G* with increasing *λ* (Fig. 3c). Values of *λ* for the base of 0.13–0.15 resulted in the exclusion of dsRNA from the Ddx4N1 condensates, in reasonable agreement with experiments.

To examine whether these results are sensitive to the simulation setup, we performed additional simulations with: (1) mixtures of 300 chains of Ddx4N1 and 15 chains of dsRNA, and (2) 400 chains of Ddx4N1 with 15 chains of dsRNA, with the elastic-network force constant tuned to account for dsRNA flexibility (see below). We did not observe substantial differences among these systems (Fig. 3c). For a given system, we note that the relationship between the bulk concentrations of Ddx4N1 and dsRNA and those in the dilute and dense phases is affected by the surface accumulation and thus by the surface-to-volume ratio of the condensates. While the thermodynamic consequence of such surface effects might depend on the relative volume of the interface and dense phase, we make the assumption that dsRNA concentration in the dilute phase is approximately constant regardless of the surface-to-volume ratio. To examine this, we compared simulations of systems containing 300 and 400 Ddx4N1 chains and found very similar Δ*G* values (Fig. 3c). Thus, for simplicity we ignore any such finite size effects. From these results, we selected the model using *λ* = 0.13 with the tuned elastic force constant, which yielded Δ*G* = 2.6 ± 0.3 kJ*/*mol, in reasonable agreement with the experimental value (2.1 ± 0.1 kJ*/*mol).

### 2.4 Elastic network tuned to capture dsRNA stiffness

In the analyses above, we treated dsRNA as a highly rigid body, but this description becomes inadequate for long chains. dsRNA has a persistence length of approximately 62 nm, ^86,87^ making it considerably stiffer than disordered ssRNA^88–90^ but comparable to dsDNA.^59^ To quantify the stiffness of the dsRNA chain, we tuned the elastic network potential to reproduce this persistence length.

We performed simulations of a single 280-bp dsRNA chain with varying elastic-network strengths (8, 10, and 20 kJ*/*mol*/*nm^2^), using the base value of *λ* = 0.13. To measure the bending stiffness, we calculated the persistence length, *l*, from the autocorrelation of displacement vectors between neighbouring base pairs,^91,92^

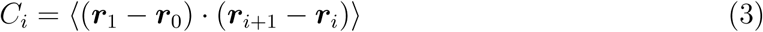

where ***r****_i_* is the centre position of the *i*-th base pair. The autocorrelation function was fitted by an exponential function (Fig. 4a),

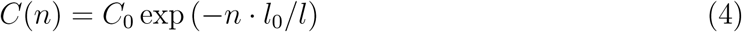

**Figure 4:**
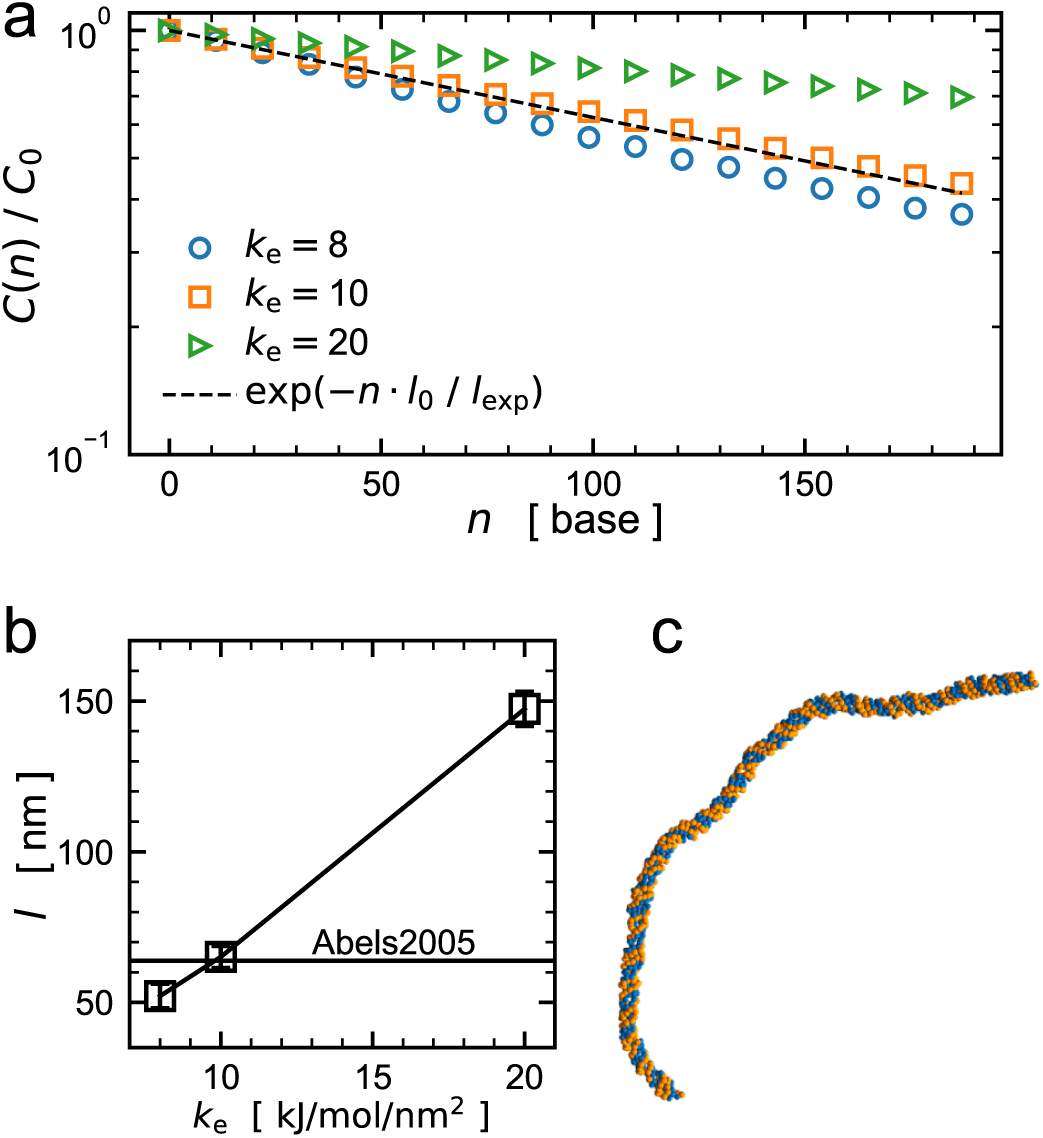
Tuning elastic-network force constant to capture dsRNA stiffness. (a) Autocorrelation function, *C*(*n*), of the base–base vectors as a function of base-pair separation, *n*, sampled every 11 base pairs in A-form dsRNA. The strength of the elastic network is indicated by labels, *k*_e_ = 8, 10, 20 kJ*/*mol*/*nm^2^. A reference curve with persistence length from Abels *et al.*^86^ is also shown, *l*_exp_ = 62 nm. (b) Persistence length, *l*, obtained by fitting an exponential model to the autocorrelation function. Error bars show standard error of the mean using time blocks. (c) A representative snapshot from simulation of *k*_e_ = 10 kJ*/*mol*/*nm^2^.

where *l* is the persistence length, *n* is the base-pair separation, and *l*_0_ is the rise per base pair. An elastic force constant of *k*_e_ = 10 kJ*/*mol*/*nm^2^ yielded a persistence length of 64 ± 2 nm (Figs. 4b and c), in agreement with the reported dsRNA persistence length (62 nm),^86^ and was therefore chosen for the CALVADOS-dsRNA model.

### 2.5 Balance between electrostatic and non-electrostatic interactions determines RNA partitioning into condensates

Based on the effect of stickiness on the dsRNA partitioning behaviour, we speculate that partitioning behaviour could be modulated by the balance between non-electrostatic interactions (governed by the AH potential) and electrostatics (governed by the Debye-Hückel, DH, potential). To explore this, we used CAPRIN1 C-terminal IDR (hereafter CAPRIN1) as a model protein, and examined ssRNA and dsRNA partitioning into CAPRIN1 condensates under different salt conditions to alter the balance between electrostatic and non-electrostatic interactions.

CAPRIN1 is a synaptic protein containing a C-terminal IDR abundant in arginine, and CAPRIN1 condensates have been shown to facilitate the dissociation of dsRNA.^20^ CAPRIN1 undergoes phase separation under high salt-screening conditions or in the presence of RNA or ATP,^93–95^ while R-to-K mutations suppress condensation.^96^ These observations indicate that condensation of the CAPRIN1 IDR is driven by both electrostatic and non-electrostatic interactions,^97^ making it a suitable model to explore both electrostatic and non-electrostatic effects.

We performed simulations of condensates formed by CAPRIN1 IDR and 12-nt ssRNA as scaffold components, using 12-bp dsRNA as a client (Figs. 5a and b). This simulation setup was designed to capture the experimental observations by Rangadurai *et al.* indicating an approximately 4,000-fold higher concentration of ssRNA than dsRNA within CAPRIN1 IDR condensates (values calculated from the reported kinetic properties).^20^ Instead of directly sampling the association and dissociation equilibrium of dsRNA, we studied the differential partitioning of ssRNA and dsRNA (Fig. 5a).^7^ In control systems, we distinguished the ssRNA client (C) chains from the scaffold (S) chains by artificially labelling them separately (ssRNA-C vs. ssRNA-S, Fig. 5b).

**Figure 5:**
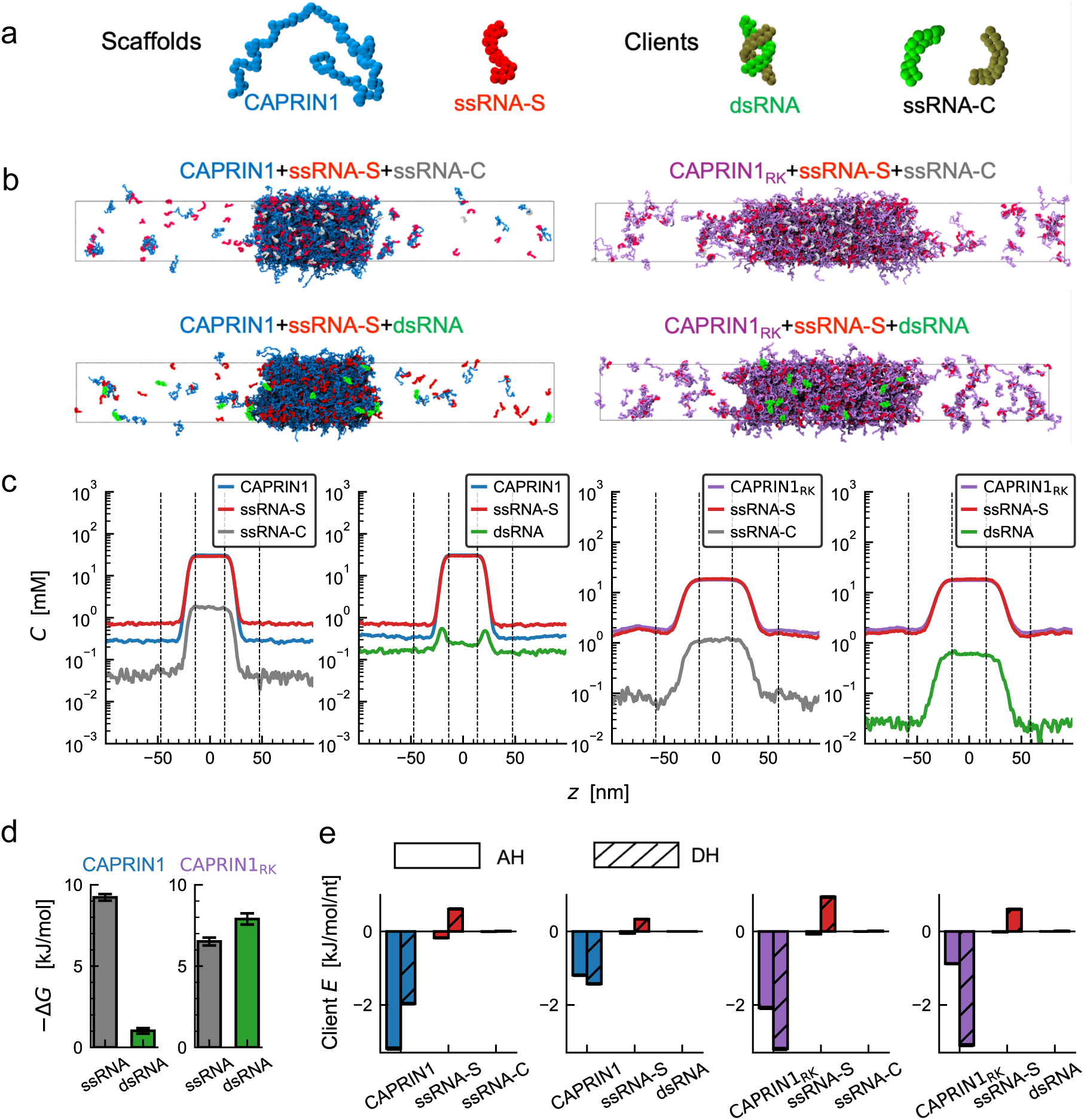
Partitioning of ssRNA and dsRNA into CAPRIN1 condensates. (a) Representative molecular structures of scaffold components (CAPRIN1 and 12-nt ssRNA) and clients (12-bp dsRNA and 12-nt ssRNA). Scaffold and client ssRNAs are denoted as ‘ssRNA-S’ and ‘ssRNA-C’, respectively, and we studied both wild-type (WT) CAPRIN1 and an R-to-K variant. (b) Representative snapshots of four simulation systems. The ionic strengths are 200 mM for WT CAPRIN1 systems and 80 mM for the variant systems. For clarity, clients are shown in the front periodic image, while scaffolds are shown in the unit cell. (c) Density profiles of scaffold components and clients for the four simulations. (d) Partitioning free energies of ssRNA and dsRNA into wild-type and R-to-K variant CAPRIN1 condensates. (e) Interaction energies between the central client chain and surrounding molecules, calculated from AH and DH potentials.

We explored salt concentrations between 80 and 200 mM and found that ssRNA-C strongly partitioned into CAPRIN1–ssRNA condensates at 200 mM salt, whereas dsRNA showed a small difference between dilute- and dense-phase concentrations and preferentially localised at the interface (Figs. 5b and c). Thus, ssRNA showed a stronger partitioning than dsRNA (Fig. 5d), indicating preferential enrichment of ssRNA within CAPRIN1 condensates.^20^ At lower salt concentrations (80 and 100 mM), dsRNA chains partitioned more strongly into the condensates than at 200 mM, and therefore showed a reduced partitioning free energy difference compared to ssRNA (Figs. S3 and S4). These results indicate that selectivity of CAPRIN1 condensates is achieved when stickiness-based interactions are sufficiently strong relative to electrostatic interactions.

To analyse the molecular interactions underlying ssRNA and dsRNA partitioning in additional detail, we calculated the interaction energies of the client RNA chain near the condensate centre with the surrounding molecules using the AH and DH potentials. ^30,69^ In all systems, client interactions with CAPRIN1 were dominant over those with ssRNA-S or with the other client chains (Figs. 5e and S5). The systems with ssRNA-C as client showed larger contributions from the AH potential relative to the DH potential at an ionic strength of 200 mM than at 100 mM, whereas dsRNA interactions were dominated by the DH potential at all salt concentrations (Figs. 5e and S5). These results demonstrate that selective ssRNA partitioning into CAPRIN1 condensates is achieved through dominant AH interactions, whereas increased DH contributions reduce the selectivity between ssRNA and dsRNA.

We subsequently investigated whether a further shift toward conditions where electrostatics are dominant would lead to loss of selectivity for ssRNA partitioning or switching to dsRNA enrichment. With this aim, we simulated condensates of the 15-R-to-K variant of CAPRIN1.^96^ The variant underwent weak phase separation at a salt concentration of 80 mM (Fig. 5b), but not at higher salt concentrations. Condensates of the CAPRIN1 variant recruited dsRNA slightly more strongly than ssRNA (Figs. 5c and d), and interactions of both clients (ssRNA-C and dsRNA) were dominated by the DH potential (Fig. 5e). These results suggest that electrostatically dominated conditions may promote selective partitioning of dsRNA over ssRNA, although the degree of selectivity we observed was small.

Collectively, our simulation results provide insight into the physical mechanism underlying selective partitioning of ssRNA and dsRNA, based on the balance between stickiness-based and electrostatic interactions. The stronger stickiness-based interactions for ssRNA explain the stronger partitioning of ssRNA compared to dsRNA in CAPRIN1 condensates (Fig. 5d). An increase in the electrostatic interactions led to weaker selectivity of ssRNA (Fig. S4). In contrast, the 15-R-to-K CAPRIN1 variant, which formed condensates at a low salt concentration (80 mM), slightly favoured partitioning of dsRNA, although this preferential enrichment has not yet been tested experimentally (Fig. 5d). We speculate that the preferential partitioning of dsRNA into condensates of 15-R-to-K CAPRIN1 is driven by entropy, as one dsRNA chain carries the same charge as two ssRNA chains, so that the release of two ssRNA chains from condensates in exchange for one dsRNA chain results in an entropic gain.^28^

### 2.6 Extension to dsDNA

Following the development of the dsRNA model, we subsequently developed a parameter set for dsDNA, which is involved, for example, in nuclear phase separation of heterochromatin and in the formation of transcriptional condensates. At the coarse-grained resolution employed here, the absence of the 2’-hydroxyl group in deoxyribose has a negligible effect on non-bonded interactions. However, this difference alters base packing and causes DNA to preferentially adopt the B-form rather than the A-form typical of dsRNA (Fig. S6a). We therefore retuned both the elastic network and the base interaction parameter *λ* for B-form dsDNA.

We tuned the elastic-network strength against experimental structural data and the base *λ* against partitioning data. An elastic-network force constant of *k*_e_ = 25 kJ*/*nm^2^*/*mol yielded persistence lengths in reasonable agreement with the experimentally reported values of approximately 45–50 nm under physiological conditions^91^ (Figs. 6a–c).

**Figure 6:**
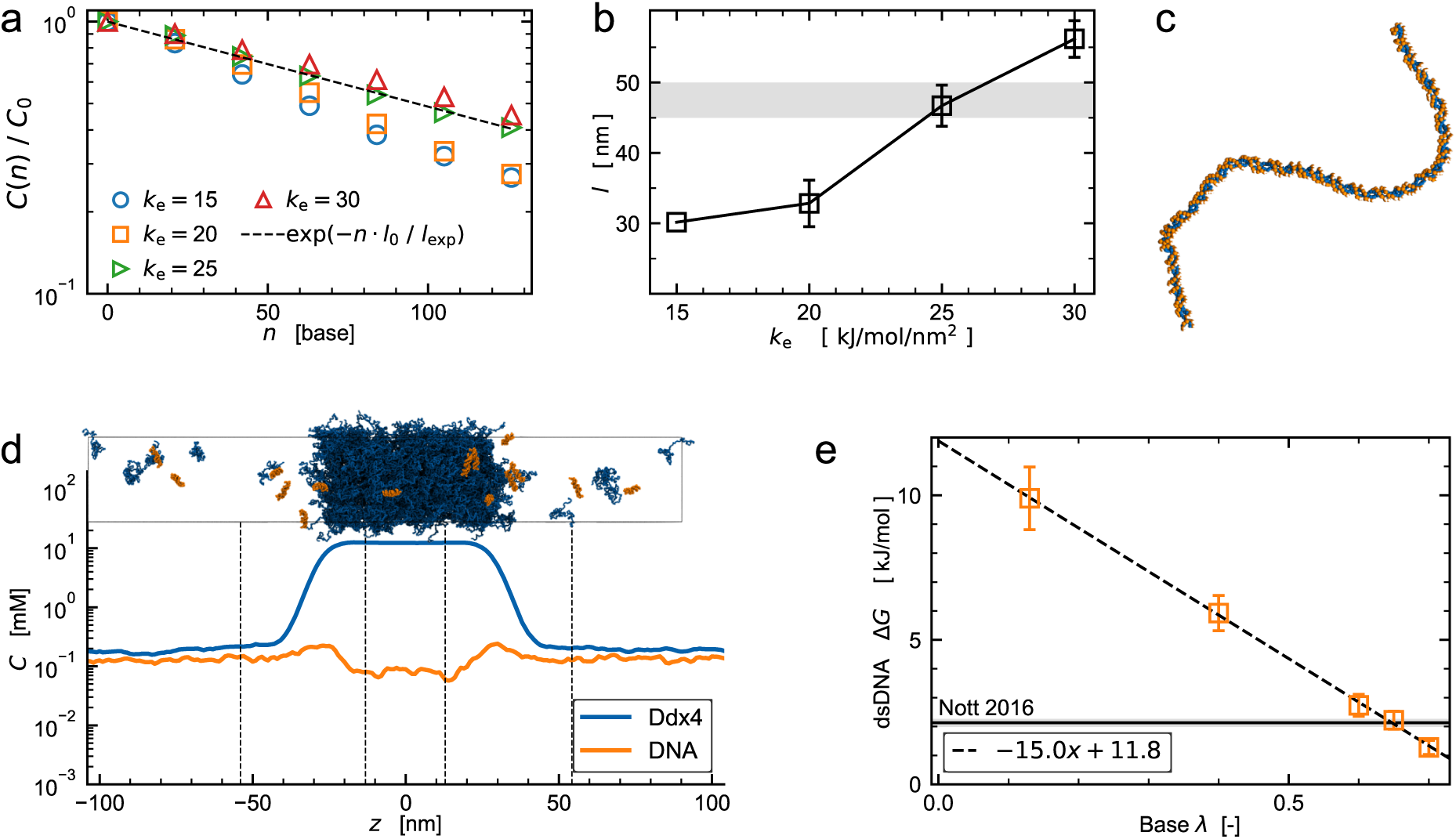
Parametrising the CALVADOS-dsDNA model. (a) Single-chain simulations of a 280-bp ACTG repeat. Autocorrelation function, *C*(*n*), of the base–base vectors as a function of base-pair separation, *n*, sampled every 11 base pairs in B-form dsDNA. The single-chain simulation was performed using base *λ* = 0.65. The strength of the elastic network is indicated by labels, *k*_e_ = 15, 20, 25, 30 kJ*/*mol*/*nm^2^. (b) Persistence length, *l*, obtained by fitting an exponential model to the autocorrelation function. Error bars show standard error of the mean using time blocks. The grey area indicates persistence length of dsDNA under physiological conditions.^91^ (c) Representative snapshot of the 280-bp ACTG repeat using *k*_e_ = 25 kJ*/*mol*/*nm^2^. Backbone and base beads are coloured in orange and blue, respectively. (d) 24-bp dsDNA partitioning into Ddx4N1 condensates. Density profiles of Ddx4N1 and dsDNA, and a representative snapshot, using base *λ* = 0.65. (e) Relationship between the base *λ* and partitioning free energy of dsDNA, Δ*G*. The black dashed line shows the linear regression model fitted to all data points. The black solid line with shaded region represents the experimental Δ*G*.

Regarding the *λ* parameters, we performed simulations of 24-bp dsDNA partitioning into Ddx4N1 condensates with varying *λ*. A value of *λ* = 0.65 yielded Δ*G* = 2.2 ± 0.3 kJ*/*mol (Figs. 6d and e), reasonably capturing weak dsDNA exclusion.^12^ However, the difference in the partitioning free energy between dsDNA and ssDNA (ΔΔ*G* = Δ*G*_ss_ − Δ*G*_ds_) was over-estimated (ΔΔ*G*_MD_ = −11.3 ± 0.5 kJ*/*mol), compared to the experimental value ΔΔ*G*_exp_ = −7.8 ± 0.7 kJ*/*mol (Tab. S1). This is because, although the experimental study reported stronger partitioning of ssRNA into Ddx4N1 condensates than ssDNA,^12^ this difference between ssDNA and ssRNA is not captured in our models.

The different stickiness parameters for the bases in dsDNA and dsRNA reflect the geometry of the B-form and A-form helices (Fig. S6a). To quantify this, we examined the relationship between *λ* and the solvent-accessible surface areas (SASA). The SASA values of nucleic acids are known to differ substantially between A- and B-form helices when a relatively large probe size is used, although they are comparable when a water-sized probe is used.^98^ In our atomistic structures, the base SASA was larger for DNA than for RNA: the sequence-averaged base SASA was 1.67 nm^2^ for RNA and 2.11 nm^2^ for DNA using the probe radius of 0.5 nm (Fig. S6b). In contrast, the coarse-grained structures did not capture this trend: the base SASA values were 0.36 nm^2^ for RNA, and 0.09 nm^2^ for DNA (Fig. S6b). This underestimation of the base exposure of dsDNA in the coarse-grained representation is compensated for by a higher base *λ*, which allows our model to reproduce the comparable partitioning behaviour observed for dsDNA.

## 3 Discussion

Previously we have parametrised the CALVADOS and CALVADOS-RNA models using data from biophysical experiments characterising single-chain properties, and reasonably predicted phase separation driven by weak multivalent interactions.^56,70^ In the CALVADOS-dsRNA and CALVADOS-dsDNA models, we calibrated the base non-electrostatic interaction parameter to modulate protein–ds nucleic acid interactions with reference to the partitioning data. Since only Ddx4N1 condensates were used for parameter tuning in this work, the general applicability of our models remains to be tested in more detail. Yet, we speculate that our models can capture weak multivalent interactions at a semi-quantitative level. Although the conditions where protein–nucleic acid condensates form vary quantitatively in a sequence-dependent manner,^28,29,99^ our models, which use common bases for all nucleotide sequences, cannot capture such sequence-dependent effects. Consequently, our models are suited for capturing nucleotide-sequence-averaged nucleic acid interactions with IDRs.

In this work, we applied our models only to duplex nucleic acids. However, we envision that they could be applied to more complex structures of mRNA and non-coding RNA by modelling ssRNA and dsRNA segments separately.^100^ The folded RNA structures could be obtained from experimentally resolved structures, or from structures computationally predicted using recent approaches.^101–103^ Additionally, amorphous structures of RNA aggregates, sampled from other coarse-grained models,^48,49,61^ could be used for simulations of RNA aggregates in protein condensates. Such applications would enable investigating how condensate environments influence the folding energy landscape of nucleic acids. ^7,8,12^

Considering that the CALVADOS models were designed to capture weak, multivalent interactions, their application to systems governed by other biophysical mechanisms requires caution. In particular, specific interactions between proteins and nucleic acids, as observed in nucleic acid–protein complexes,^24,26,104,105^ are not fully captured unless the model is augmented with restraint potentials that encode such specificity. Similarly, intermolecular base-pairing interactions are not explicitly described, and thus the model is not expected to capture RNA condensation driven by base pairing.^48,49,61^ Furthermore, our dsRNA and dsDNA models restrain nucleic acid conformation to the initial structures rather than sampling folding–unfolding equilibria. These aspects may be addressed in future extensions of the model.

## 4 Conclusions

Here, we present a coarse-grained model for dsRNA, building on our two-bead-per-nucleotide model of ssRNA.^56^ This extension included (i) an elastic network potential to maintain the helical conformation, and (ii) a reduced stickiness assigned to the base bead, reflecting that base atoms of dsRNA are less accessible to the surrounding molecules than those of ssRNA. We calibrated the force constant for the polymer stiffness of dsRNA using measurements of the persistence length. We parametrised the stickiness parameter for the bases using measurements of dsRNA–Ddx4N1 interactions probed by the partitioning free energy into Ddx4N1 condensates. We also used this procedure to develop a dsDNA model. Our models capture both overall conformations of double-stranded nucleic acids and their interactions with IDRs, enabling simulation of mixed double-stranded nucleic acid–protein condensates. Furthermore, we demonstrated the role of electrostatic and stickiness-based interactions in selective dsRNA and ssRNA partitioning into CAPRIN1 condensates. We envision that our models will contribute to a better understanding of how structures of RNA, via chemistry-specific interactions, help condensates recognise specific types of RNA, and drive mixing and demixing with protein condensates.

## 5 Methods

### 5.1 Force field of CALVADOS-dsRNA and -dsDNA models

The potential energy of the two-bead-per-nucleotide model is expressed as

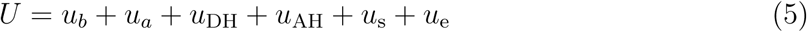

where *u_b_*, *u_a_*, *u*_DH_, *u*_AH_, *u*_s_, *u*_e_ are the bond, angle, electrostatic, short-ranged stickiness, neighbouring stacking and elastic network terms, respectively. The bond potential is applied to backbone–backbone and backbone–base distances, and is described by a harmonic potential,

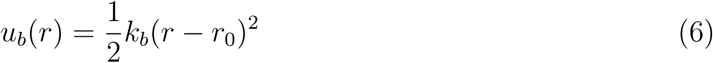

where *k_b_* and *r*_0_ are the force constant and equilibrium bond length, respectively, and with *k_b_* = 1400 kJ*/*mol*/*nm^2^ for the backbone–backbone bond, and 2200 kJ*/*mol*/*nm^2^ for the backbone–base bond. *r*_0_ is calculated from the initial structure of the dsRNA or dsDNA. The angle potential, accounting for the stiffness of the backbone chain, is described by a harmonic potential,

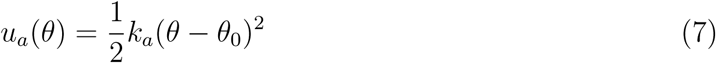

where *k_a_* is the spring constant determined for the backbone chain to be 4.2 kJ*/*mol*/*rad^2^. *θ*_0_ is the equilibrium angle, calculated from the initial structure of dsRNA or dsDNA. The electrostatic interaction is described by the DH potential,

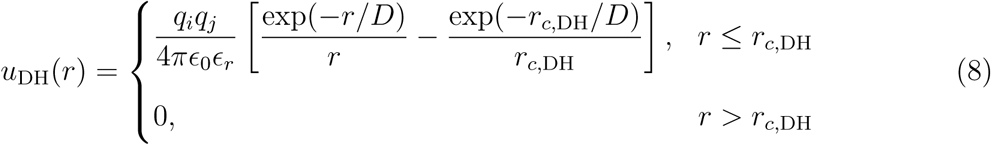

where *q* is the charge of the beads, i.e. −*e* for the backbone bead and 0 for the base, *ɛ*_0_ is the vacuum permittivity, *ɛ_r_* is the dielectric constant of water with an empirical temperature dependence,^106^

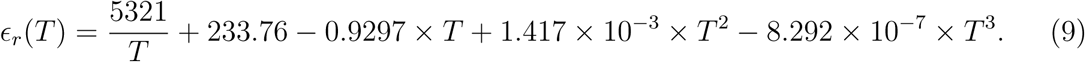

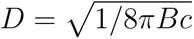 is the Debye length of an electrolyte solution of ionic strength, *c*, and *B*(*ɛ_r_*) is the Bjerrum length as a function of *ɛ_r_*(*T*). *r_c,_*_DH_ is the cutoff distance for the DH potential, set to 4 nm. The short-range interaction is described by the AH potential,^107^

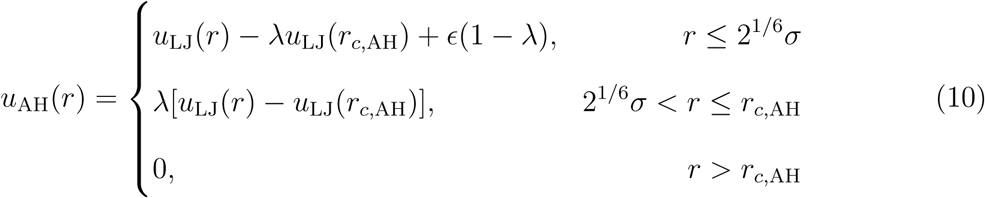

where *u*_LJ_(*r*) is the LJ potential with *ɛ*=0.8368 kJ mol*^−^*^1^, and the diameters, *σ*, are 0.6954 nm and 0.6238 nm for the backbone and the base, respectively. The cutoff distance for the AH potential, *r_c,_*_AH_, was set to 2 nm. Both AH and DH interactions were applied to protein– protein, protein–nucleic acid, and nucleic acid–nucleic acid interactions in the same manner. For the AH potential between different types of beads, combination rules were applied, i.e., *σ* = (*σ_i_*+*σ_j_*)*/*2 and *λ* = (*λ_i_*+*λ_j_*)*/*2. Any pair of beads subject to *u_b_*, *u_a_*, *u_s_* or *u_e_* was excluded from *u*_AH_ and *u*_DH_. The interaction between the neighbouring base beads is described by a stacking potential of the following functional form,^56^

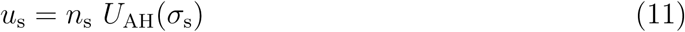

where *n*_s_ is a scaling factor. *U*_AH_(*σ*_s_) is the AH potential using the *λ* of the base, with *σ*_s_ = *d/*(2^1^*^/^*^6^) being the diameter parameter calculated from the distance, *d*, in the initial structure. We used *n*_s_ = 15 at *λ* = 1.18 for ssRNA, whereas for dsRNA and dsDNA we set *n*_s_ = 136 to compensate for their lower base *λ* values. The elastic network term is described by a harmonic potential,

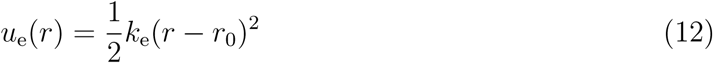

where *k*_e_ is the spring force constant and *r*_0_ is the equilibrium distance in the initial structure. The elastic network was applied to bead pairs within 1.5 nm, corresponding approximately to interactions extending to ±2 neighbouring nucleotides for the A-form dsRNA. *k*_e_ values for dsRNA and dsDNA were 10 kJ*/*mol*/*nm^2^ and 25 kJ*/*mol*/*nm^2^, respectively. The elastic network potential was not applied to pairs of beads subject to any of *u_b_*, *u_a_*, or *u*_s_.

### 5.2 Setup for systems with IDRs and nucleic acids

The Ddx4N1 sequence, the N-terminal region of Ddx4 (UniProt ID: Q9NQI0, Residues 1– 245) fused to a five-residue N-terminal extension (GAMGS) for purification,^108^ was taken from the work by Nott *et al.* ^12^ (Tab. S2). ssRNA contained a generic (non-sequence-specific) 24-nt sequence, which we modelled as a flexible chain without local secondary structures.^56^ The systems of Ddx4N1 with either ssRNA, dsRNA or dsDNA used an ionic strength of 150 mM, a temperature of 293 K, and pH 8 to determine the charge of histidine residues. For Ddx4N1–ssRNA systems, we used a simulation box with a box size of (30 nm, 30 nm, 200 nm) containing 400 Ddx4N1 chains and 16–32 ssRNA chains (Tab. S3). For low RNA ratio systems, we used a system with (50 nm, 50 nm, 200 nm) containing 800 Ddx4N1 chains and 16 ssRNA chains, and a system with (40 nm, 40 nm, 200 nm) containing 800 Ddx4N1 chains and 8 ssRNA chains.

For Ddx4N1–dsRNA systems, we used a simulation box size of (30 nm, 30 nm, 210 nm), containing 300 or 400 chains of Ddx4N1 and 15 chains of dsRNA (Tab. S3). The dsRNA sequence consists of six repeats of ACUG, and was modelled as an A-form helix using a builder^109^ (https://github.com/sbottaro/build_aform), modified here to include the 5’-terminal phosphate. From the atomistic helical model, we obtained the coarse-grained geometry by placing the backbone and base beads at the phosphorus atom and the nitrogen atom (N9 for Ade and Gua, and N1 for Cyt and Ura), respectively. We assigned base *λ* values in the range of 0–0.3, and employed a constraint with either a strong elastic force constant (*k*_e_ = 7000 kJ*/*mol*/*nm^2^) or the calibrated elastic force constant. For the strongly constrained dsRNA model, the stacking potential changed according to the chosen value of *λ* and a constant (*n_s_* = 15), but this effect was negligible because the duplex structure was maintained mainly by the elastic network. For the flexibility-tuned model, we used the corrected value (*n_s_* = 136).

Similarly, the sequence of dsDNA is six repeats of ACTG, and its coarse-grained geometry was obtained from an atomistic B-form dsDNA structure (Tab. S2). The dsDNA structure was built using MDNA,^110^ with modifications made to include the 5’-terminal phosphate. We used the base *λ* values in the range of 0.13–0.70, and a constraint with the calibrated elastic force constant. Ddx4N1–dsDNA systems used the same box geometry as the Ddx4N1– dsRNA systems, and contained 400 Ddx4N1 chains and 15 dsDNA chains.

The CAPRIN1 sequence, the C-terminal region (UniProt ID: Q14444, Residues 607–709) with the N623TN630T double mutation, and the 12-nt RNA sequences were taken from the work by Rangadurai *et al.*^20^ (Tab. S2). The 12-bp dsRNA structure was built similarly to the 24-bp dsRNA. The sequences of client and scaffold ssRNAs were identical. The R-to-K variant was generated by substituting all 15 arginine residues with lysine in the double-mutant variant of CAPRIN1. We used a simulation box size of (25 nm, 25 nm, 200 nm) containing 500 chains of CAPRIN1 (or its R-to-K variant), 500 chains of scaffold ssRNA, and client RNA (15 chains of dsRNA or 30 chains of ssRNA). We set ionic strengths in the range of 80–200 mM, a temperature of 293 K, and pH 7.

All proteins were modelled by the CALVADOS 2 force field.^70^ Parameters for ssRNA,^56^ dsRNA, and dsDNA are shown in Tab. 1. All simulation input files were generated using the CALVADOS package.^111^

**Table 1:**
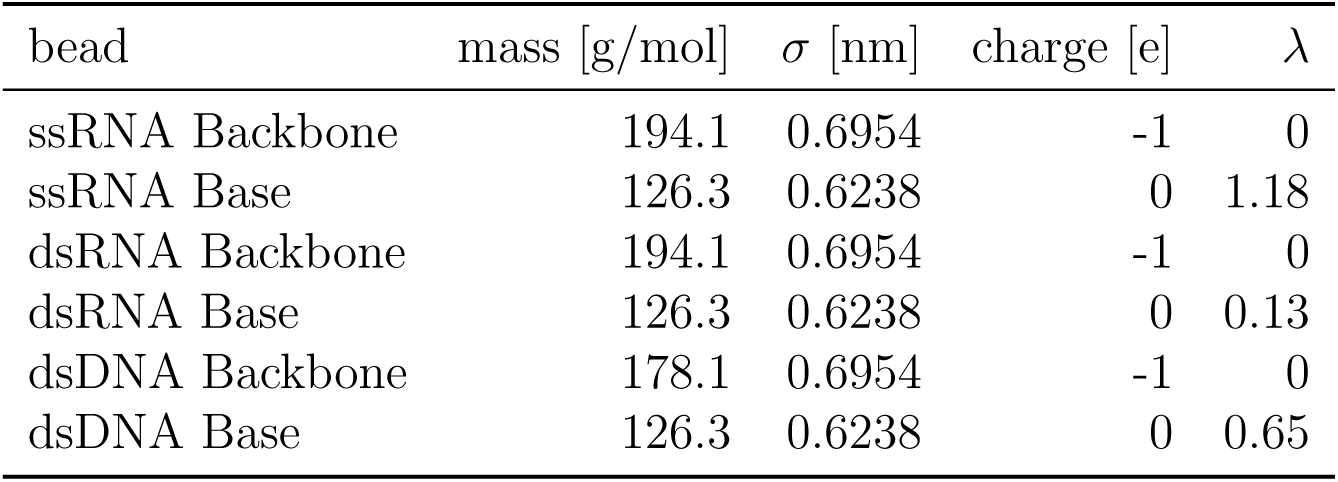
Parameters of ssRNA, dsRNA and dsDNA (mass, molecular diameter and charge of beads, stickiness).

### 5.3 Molecular dynamics simulations of IDR condensates with nucleic acids

All simulations were performed using OpenMM 8.2.0.^112^ We used a Langevin integrator with a time step of 10 fs and a friction coefficient of 0.01 ps*^−^*^1^. The initial configurations were generated by randomly placing proteins in a cubic arrangement, ssRNA in an Archimedean spiral, and dsRNA in the modelled structures. Biased simulations were performed, with an external force dragging the components toward the centre of the box along the *z*-axis, to form a single slab at the centre of the box. Subsequently, unbiased simulations were performed for at least 5.8 *µ*s (Tab. S3). Trajectories were output every 10 ns, and the first 1 *µ*s of each simulation was removed from analysis. The convergence of the simulations was checked based on the cumulative mean and estimated standard error of the mean using the time-blocking method^113^ as implemented in https://github.com/fpesceKU/BLOCKING (Figs. S7, S8, S9, and S10).

### 5.4 Partitioning free energy

To remove the drift of condensates, we calculated the density profile of scaffold IDRs for each time step, identified the centre of the condensate, and translated the condensate to the centre of the simulation box. Subsequently, density profiles for each component were computed along the longest box axis using a bin width of 1 nm. We determined the position of the dense phase, dilute phase, and interface by fitting the semi-profiles of the scaffold IDR at *z <* 0 and 0 *< z* to the function of the interfacial density curve,

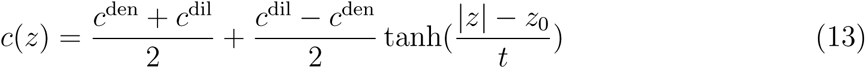

where *c*^den^, *c*^dil^, *t*, *z*_0_ are fitting parameters. The dense phase region was defined as |*z*| *< z*_0_ − *β*_den_*t* and the dilute phase as |*z*| *> z*_0_ + *β*_dil_*t*, with *β*_den_ and *β*_dil_ calibrated for each system. We averaged the density of the two phases over the defined region for each time frame, and obtained concentrations of the phases. The standard errors of the means of *c*^den^ and *c*^dil^ were estimated using the time-blocking approach. The partitioning free energy was defined as

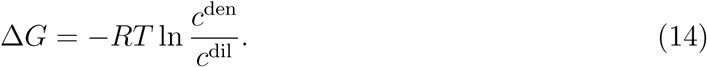

The standard error of Δ*G* was calculated using error propagation from those of *c*^den^ and *c*^dil^. For the Ddx4N1–ssRNA systems with low RNA concentrations, the standard error was calculated from multiple replicas.

### 5.5 Interaction energy analysis for CAPRIN1 systems

We selected chain *A* of molecular type *α* (client RNA) located at the centre of the box, and computed the per-nucleotide interaction energy between the central client and molecular type *β*, separately for AH and DH potentials,

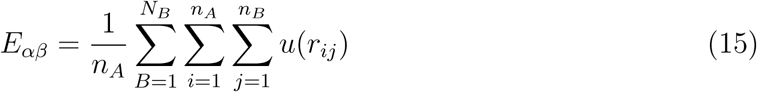

where *u* is *u*_AH_ or *u*_DH_, *N*_B_ is the number of chains of molecular type *β*, *n_A_* (*n_B_*) is the sequence length of chain *A* (*B*), *B* is the chain index for molecule *β*, and *i* (*j*) runs over residues of chain *A* (*B*). *α* is client RNA (ssRNA-C or dsRNA), and *β* is either CAPRIN1, scaffold ssRNA, or non-central client. All chains of molecular type *β* were included in the sum, although due to the interaction cutoff, only particles *j* within 4 nm of any particle *i* in chain *A* contributed. The per-nucleotide interaction energy of chain A was calculated for a direct comparison between ssRNA and dsRNA. The central chain was re-identified at each time point, and the mean *E_αβ_* over time was calculated. The standard error was estimated using the time-blocking approach.

### 5.6 Single-chain simulation of dsDNA and dsRNA

The sequences of dsRNA and dsDNA were 70 repeats of ACUG and ACTG, respectively, with their complementary strands. Coarse-grained structures of dsRNA and dsDNA were built from their helical models, similarly to the 24-bp dsRNA and 24-bp dsDNA. We assigned elastic force constant values of 8, 10, and 20 kJ*/*mol*/*nm^2^ for dsRNA, and 15, 20, 25, and 30 kJ*/*mol*/*nm^2^ for dsDNA. Single-chain 280-bp dsRNA (or dsDNA) was placed in a cubic box with an edge length of 150 nm. All simulations were performed at a temperature of 293 K, and a salt concentration of 150 mM, and continued for 2.0 *µ*s with trajectories output every 1 ns.

### 5.7 Persistence length

The persistence length, a metric of stiffness, has been used to characterise structural ensembles from molecular simulations of both dsRNA and dsDNA.^38,59,60^ We calculated the persistence length using the autocorrelation function (Eq. 3), and the persistence length was obtained as a fitting parameter (Eq. 4). In the 2.0 *µ*s simulation, the first 0.1 *µ*s of the trajectory was removed from analysis. The standard error of the mean was estimated using the time-blocking approach, where the 1.9 *µ*s trajectory was divided into 0.38 *µ*s blocks.

## Acknowledgement

I.Y. acknowledges support by JSPS through Overseas Research Fellowships, and JST through ACT-X (JPMJAX24LJ, to I.Y.). E.Y. acknowledges support from JST (PRESTO, JPMJPR22EE, to E.Y.). The research was also supported by the PRISM (Protein Interactions and Stability in Medicine and Genomics) centre, funded by the Novo Nordisk Foundation (NNF18OC0033950, to K.L.-L.). We acknowledge access to computational resources from the ROBUST Resource for Biomolecular Simulations (supported by the Novo Nordisk Foundation grant no. NF18OC0032608).

## 6 Code Availability

The code and parameters for the CALVADOS-dsRNA *β*.0 model are available at the CALVA-DOS GitHub page at https://github.com/KULL-Centre/CALVADOS. Additional data and scripts for this paper are available at https://github.com/KULL-Centre/_2026_yasuda_dsNA.

## 8 Supporting information

**Table S1:** Partitioning free energy, Δ*G*, of RNAs and DNAs into Ddx4N1 condensates under conditions corresponding to the measurements by Nott *et al.*^12^ ΔΔ*G* = Δ*G*_ss_ − Δ*G*_ds_ represents the difference between ss- and ds-forms. All energies are in kJ/mol. *: Taken from the ssRNA result. Because the only difference is the mass of the backbone bead, the same thermodynamic properties are expected.

| Client | Exp $\Delta G$ [kJ/mol] | MD $\Delta G$ [kJ/mol] | Exp $\Delta\Delta G$ [kJ/mol] | MD $\Delta\Delta G$ [kJ/mol] |
| --- | --- | --- | --- | --- |
| 24nt ssRNA | $-9.1 \pm 0.4$ | $-9.1 \pm 0.4$ | $-11.2 \pm 0.4$ | $-11.7 \pm 0.5$ |
| 24bp dsRNA | $2.1 \pm 0.1$ | $2.6 \pm 0.3$ | | |
| 24nt ssDNA | $-5.9 \pm 0.6$ | $-9.1 \pm 0.4^*$ | $-7.8 \pm 0.7$ | $-11.3 \pm 0.5$ |
| 24bp dsDNA | $1.9 \pm 0.4$ | $2.2 \pm 0.3$ | | |

**Table S2:**
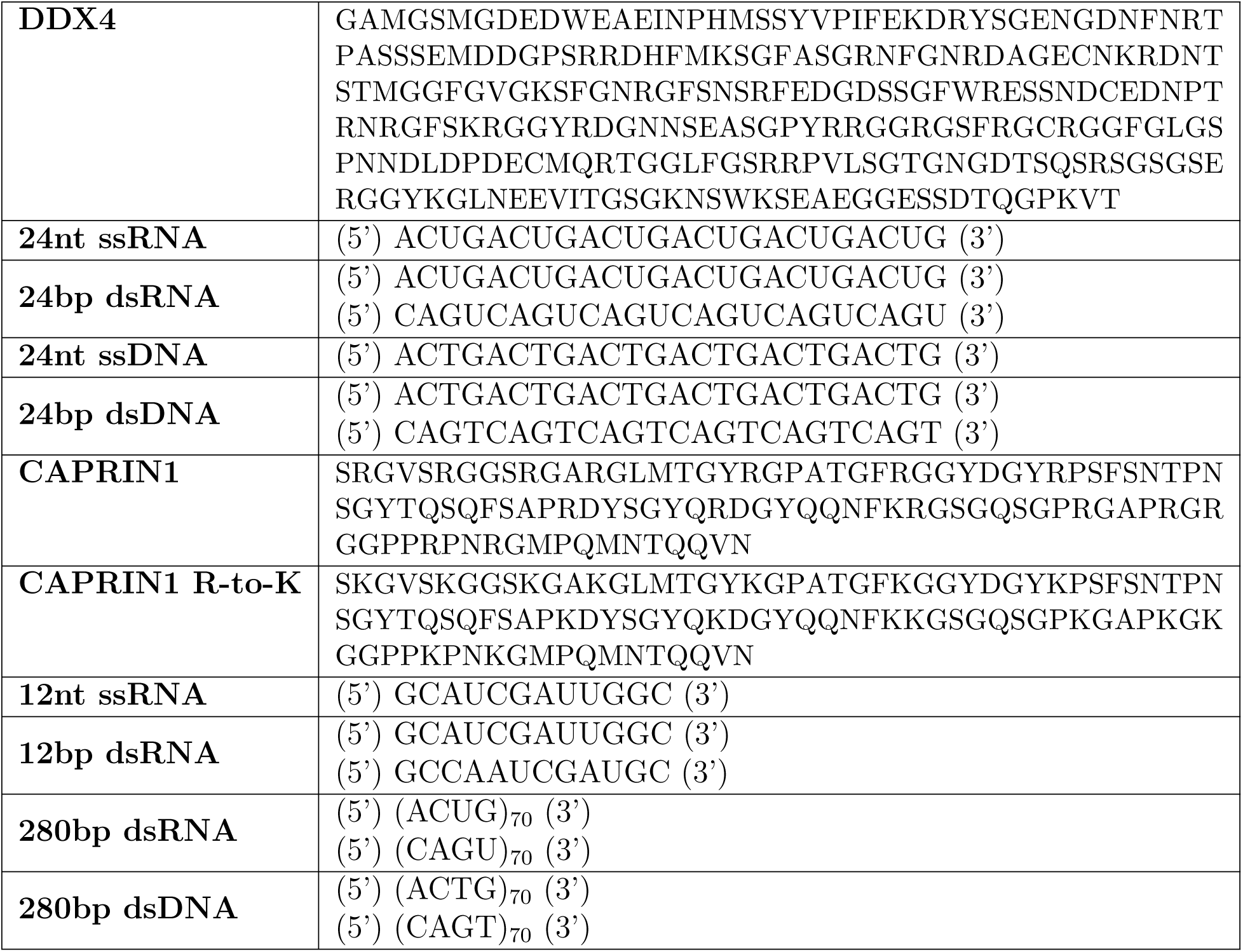
Sequences used in this work.

**Table S3:**
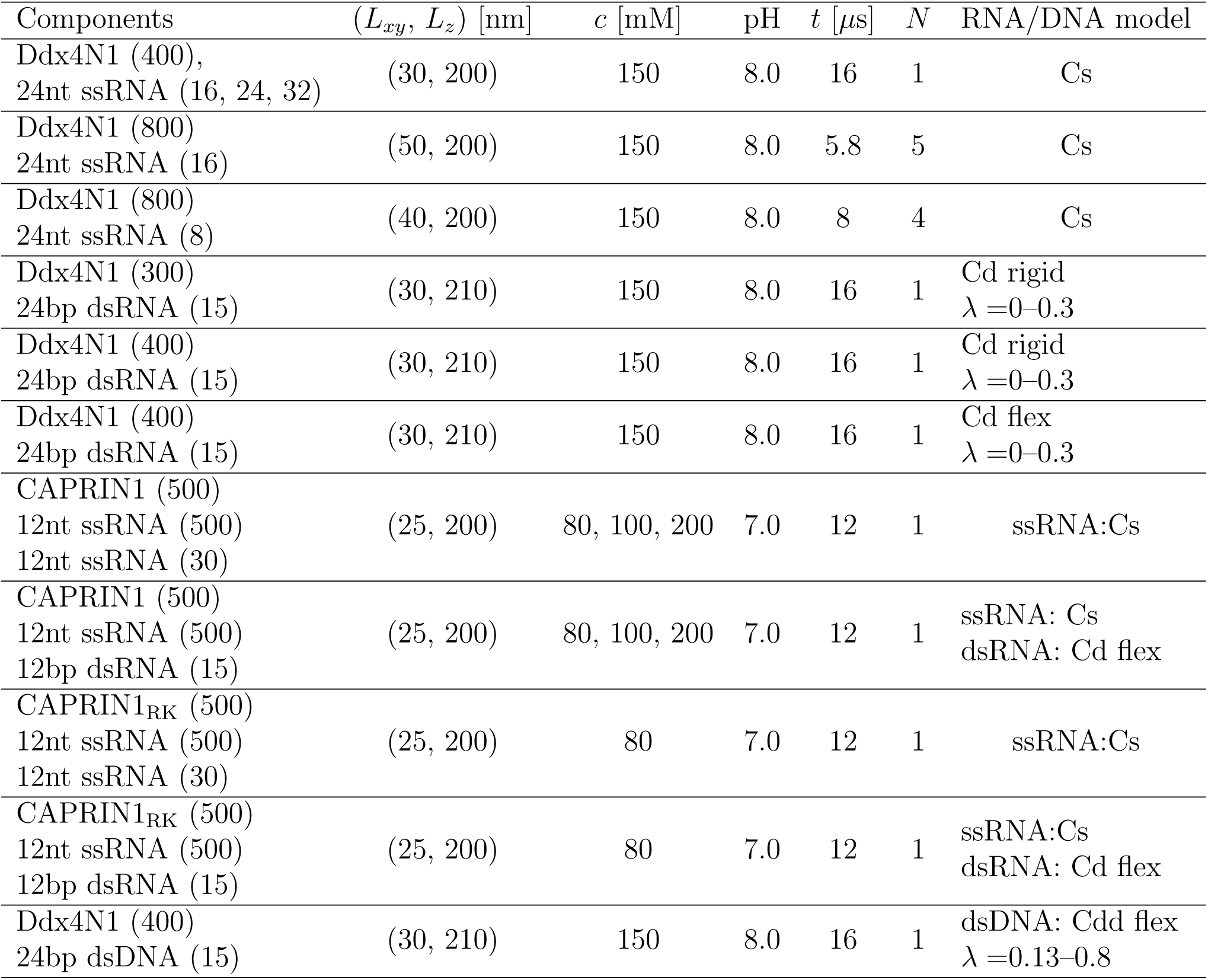
Simulation systems. System geometry (*L_xy_*, *L_z_*), ionic strength (*c*), simulation length (*t*), and number of replicas (*N*). RNA model abbreviations: CALVADOS-ssRNA (Cs), CALVADOS-dsRNA (Cd) and CALVADOS-dsDNA (Cdd). Elastic networks with strong and optimised spring constants are denoted as ‘rigid’ and ‘flex’, respectively. Stickiness of base (*λ*). The temperature of all simulations was set to 293 K.

| Components | $(L_{xy}, L_z)$ [nm] | $c$ [mM] | pH | $t$ [μs] | $N$ | RNA/DNA model |
| --- | --- | --- | --- | --- | --- | --- |
| Ddx4N1 (400),<br>24nt ssRNA (16, 24, 32) | (30, 200) | 150 | 8.0 | 16 | 1 | Cs |
| Ddx4N1 (800)<br>24nt ssRNA (16) | (50, 200) | 150 | 8.0 | 5.8 | 5 | Cs |
| Ddx4N1 (800)<br>24nt ssRNA (8) | (40, 200) | 150 | 8.0 | 8 | 4 | Cs |
| Ddx4N1 (300)<br>24bp dsRNA (15) | (30, 210) | 150 | 8.0 | 16 | 1 | Cd rigid<br>$\lambda = 0-0.3$ |
| Ddx4N1 (400)<br>24bp dsRNA (15) | (30, 210) | 150 | 8.0 | 16 | 1 | Cd rigid<br>$\lambda = 0-0.3$ |
| Ddx4N1 (400)<br>24bp dsRNA (15) | (30, 210) | 150 | 8.0 | 16 | 1 | Cd flex<br>$\lambda = 0-0.3$ |
| CAPRIN1 (500)<br>12nt ssRNA (500)<br>12nt ssRNA (30) | (25, 200) | 80, 100, 200 | 7.0 | 12 | 1 | ssRNA:Cs |
| CAPRIN1 (500)<br>12nt ssRNA (500)<br>12bp dsRNA (15) | (25, 200) | 80, 100, 200 | 7.0 | 12 | 1 | ssRNA: Cs<br>dsRNA: Cd flex |
| CAPRIN1<sub>RK</sub> (500)<br>12nt ssRNA (500)<br>12nt ssRNA (30) | (25, 200) | 80 | 7.0 | 12 | 1 | ssRNA:Cs |
| CAPRIN1<sub>RK</sub> (500)<br>12nt ssRNA (500)<br>12bp dsRNA (15) | (25, 200) | 80 | 7.0 | 12 | 1 | ssRNA:Cs<br>dsRNA: Cd flex |
| Ddx4N1 (400)<br>24bp dsDNA (15) | (30, 210) | 150 | 8.0 | 16 | 1 | dsDNA: Cdd flex<br>$\lambda = 0.13-0.8$ |

**Figure S1:**
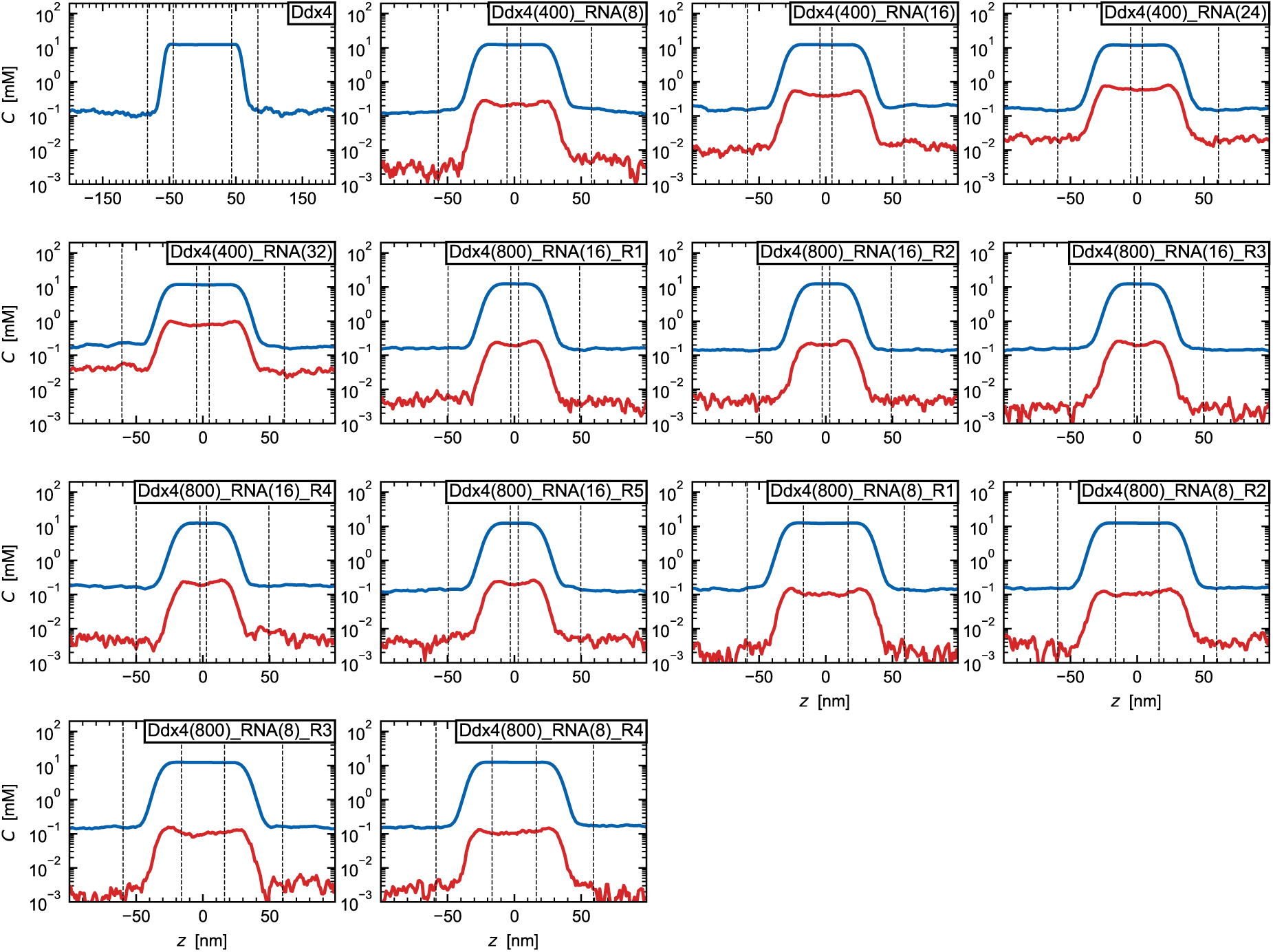
Density profiles of Ddx4N1–ssRNA systems. Ddx4N1 and ssRNA are shown in blue and red, respectively.

**Figure S2:**
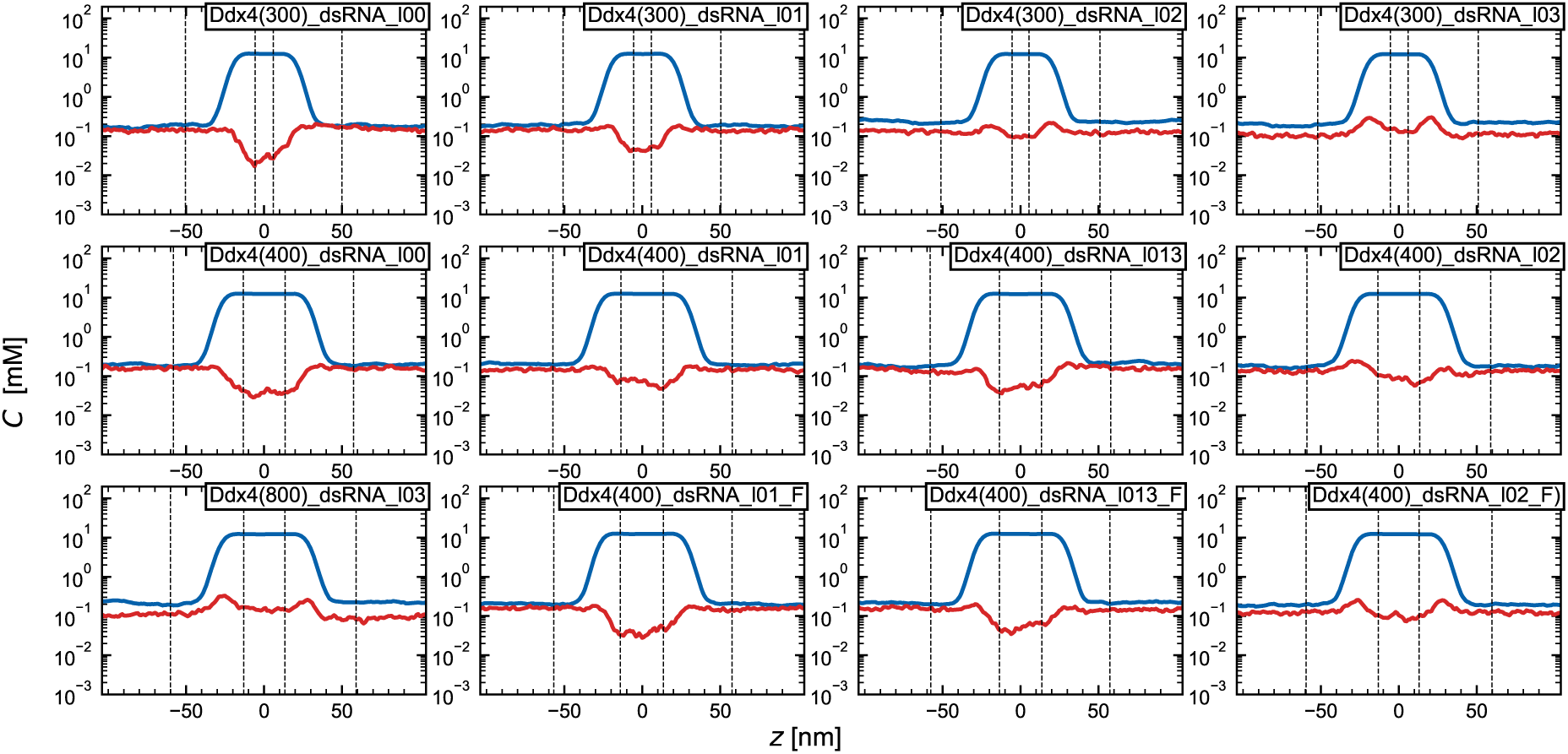
Density profiles of Ddx4N1–dsRNA systems. Ddx4N1 and dsRNA are shown in blue and red, respectively.

**Figure S3:**
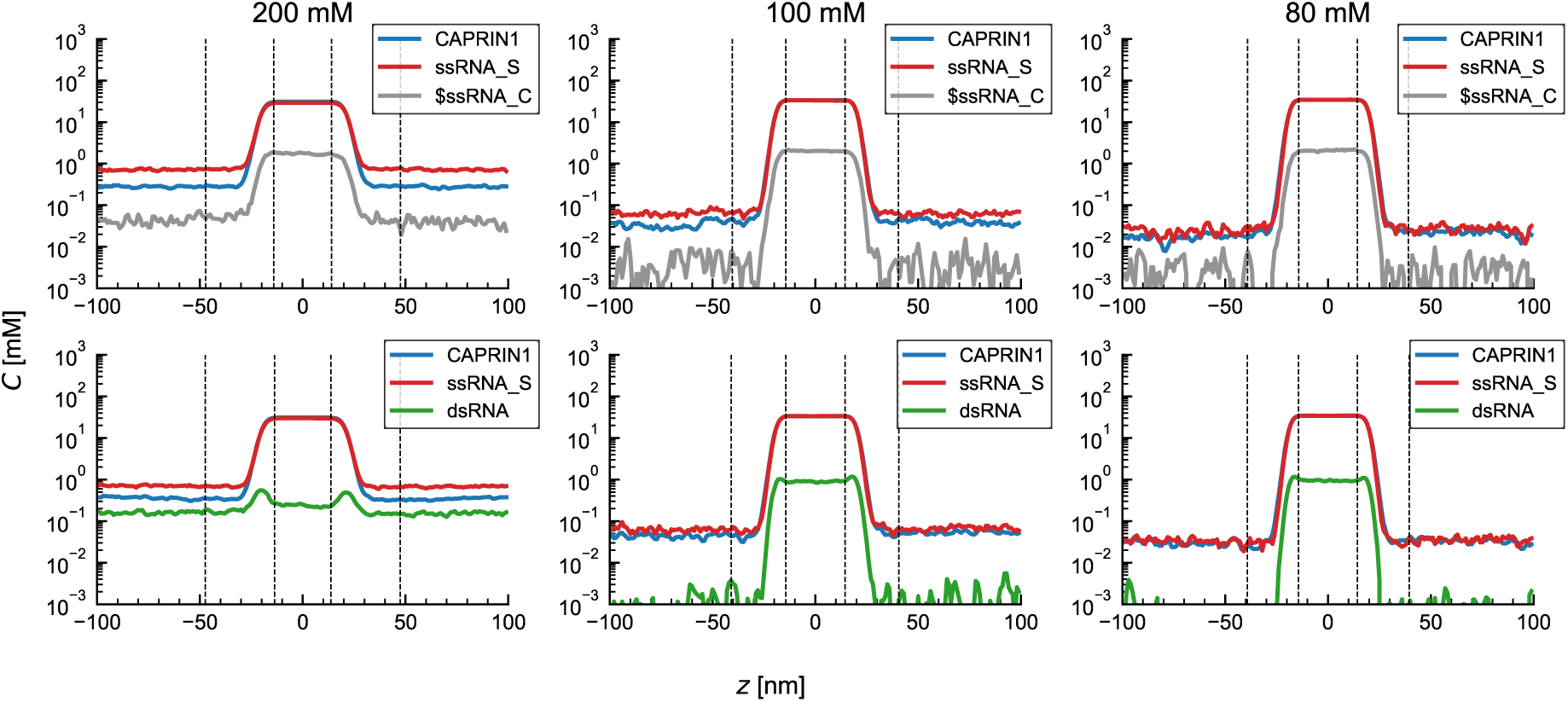
Density profiles of CAPRIN1–ssRNA–client (ssRNA-C or dsRNA) systems at salt concentrations of 80, 100 and 200 mM.

**Figure S4:**
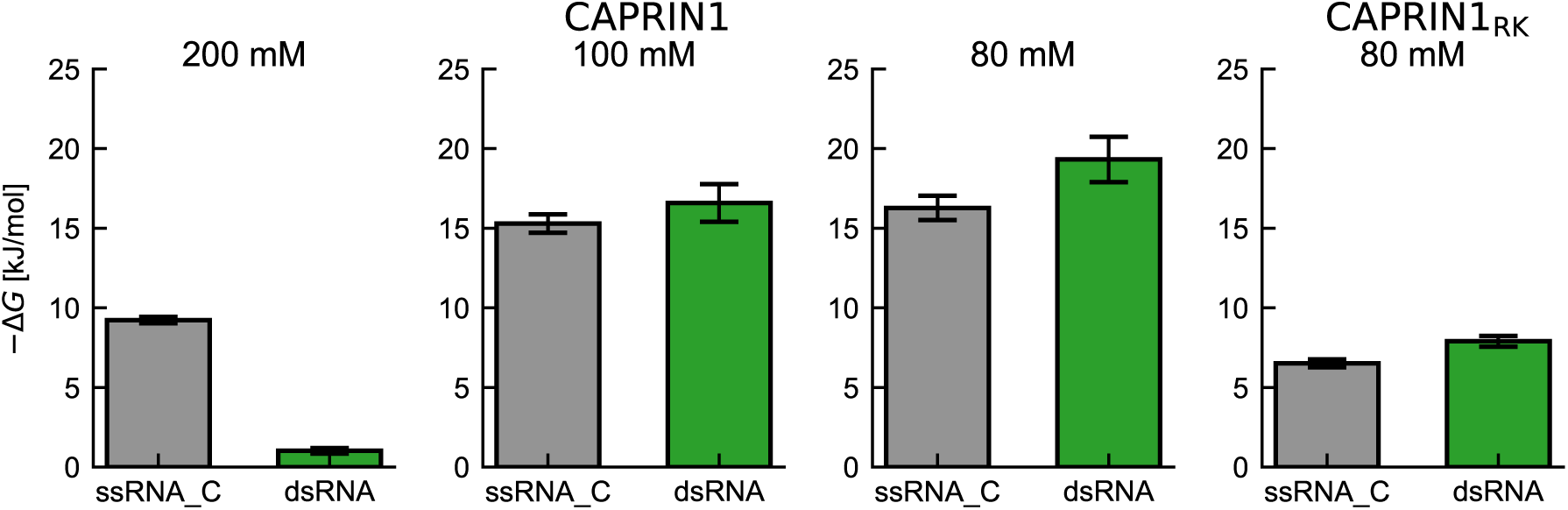
Partitioning free energy into CAPRIN1–ssRNA condensates. Systems shown: CAPRIN1–ssRNA–client at 80, 100 and 200 mM salt concentrations; CAPRIN1_RK_–ssRNA– client system at 80 mM.

**Figure S5:**
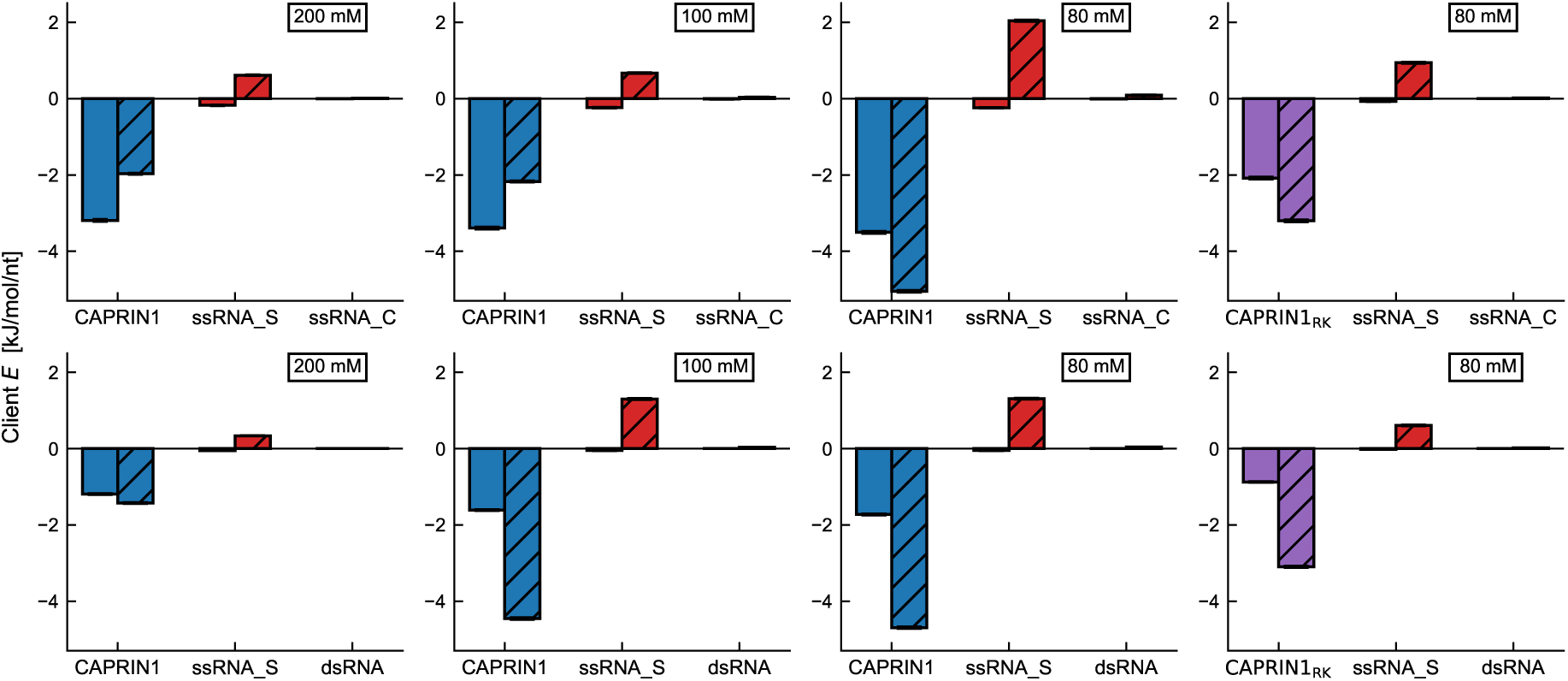
Interaction energies between the central client chain and surrounding molecules, calculated from AH and DH potentials. Systems shown: CAPRIN1–ssRNA–client at 80, 100 and 200 mM salt concentrations; CAPRIN1_RK_–ssRNA–client system at 80 mM.

**Figure S6:**
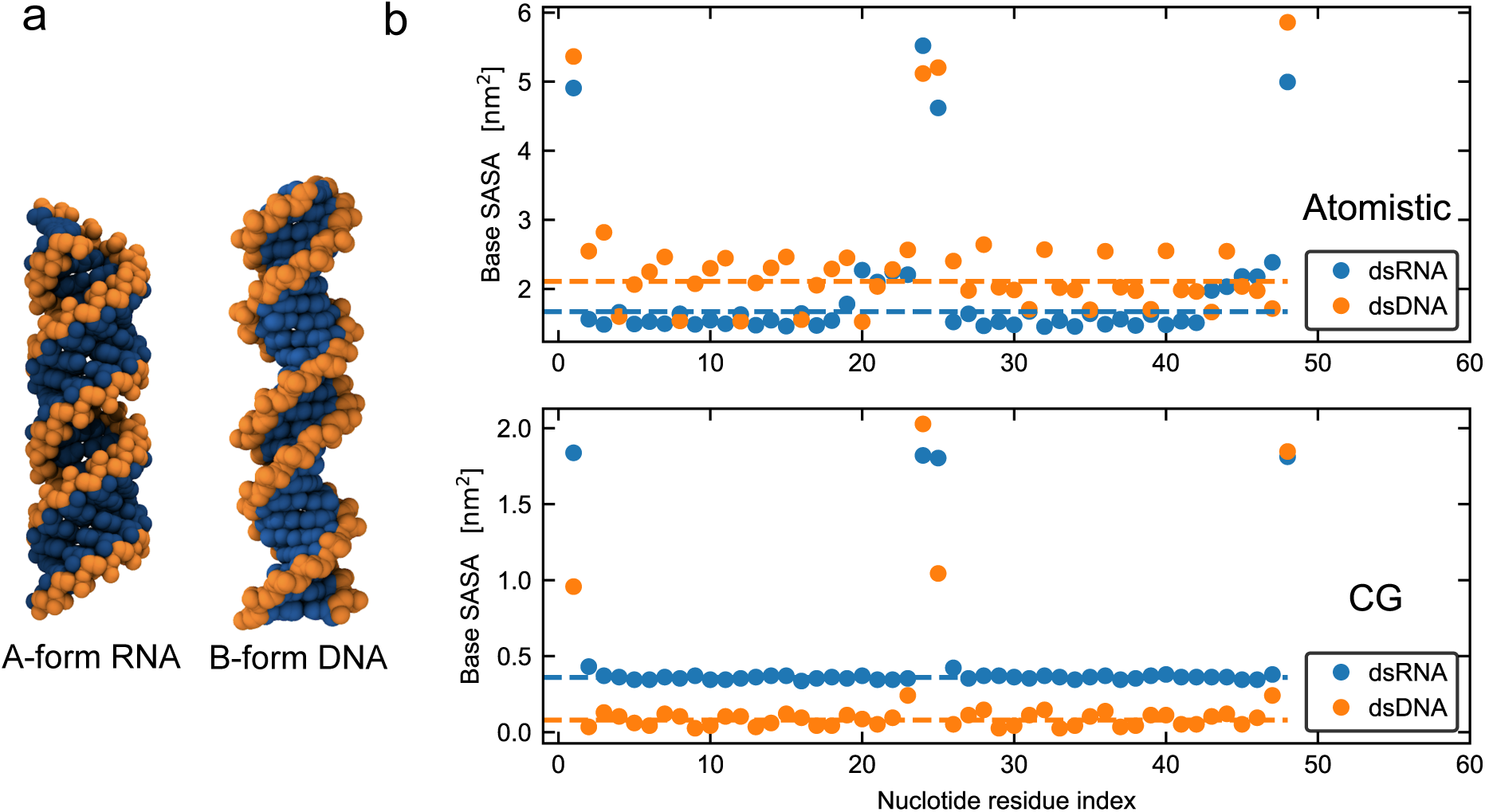
Solvent-accessible surface area (SASA) of 24-bp DNA and RNA. (a) Atomistic representation of A-form dsRNA and B-form dsDNA. Only heavy atoms were used for visualisation. (b) SASA of bases per nucleotide, calculated using a probe radius of 0.5 nm. For the atomistic structures, the base SASA was computed as the sum of the SASA of the heavy atoms in the bases. For the coarse-grained structures, the SASA of the base beads was computed. Dashed lines indicate the sequence-averaged SASA values. The terminal nucleotides were excluded from the calculation of the sequence-averaged SASA values.

**Figure S7:**
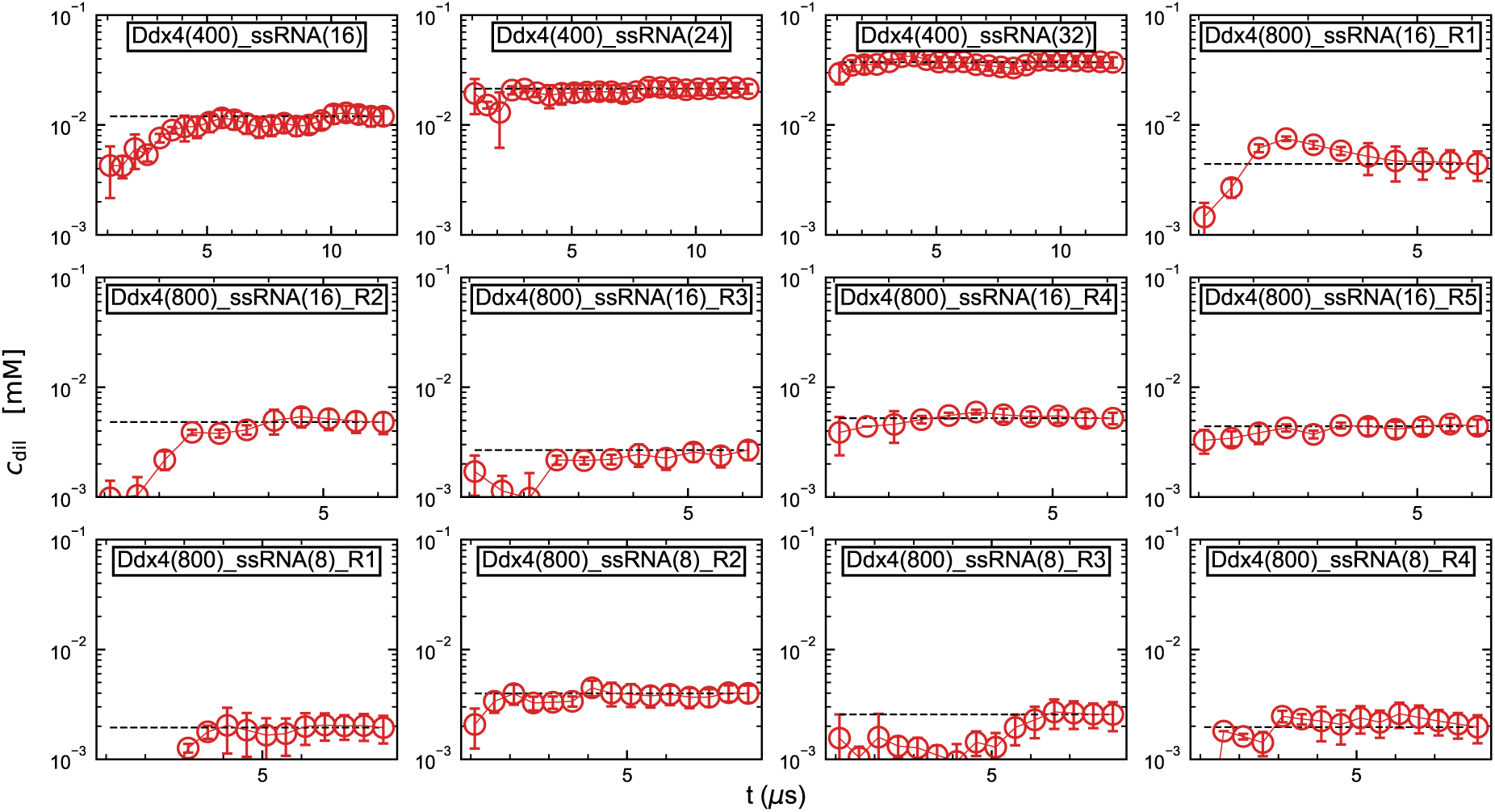
Convergence assessment of the Ddx4N1–ssRNA systems based on the dilute-phase concentration of ssRNA, *c*_dil_. Error bars show the standard error of the mean. The number in parentheses indicates the number of chains of each component, and ‘R’ denotes the replica number.

**Figure S8:**
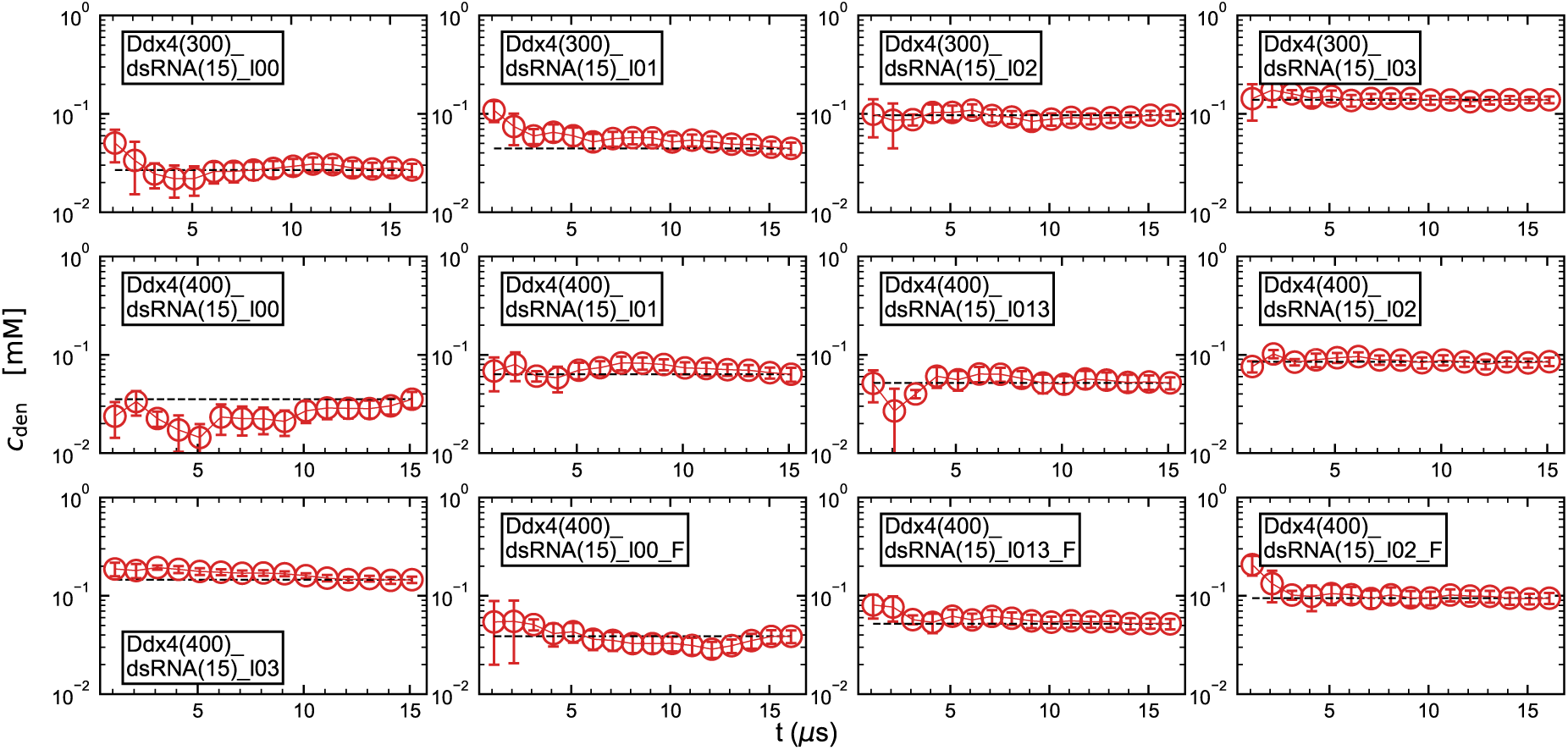
Convergence assessment of the Ddx4N1–dsRNA systems based on the dense-phase concentration of dsRNA. Error bars show the standard error of the mean. The number in parentheses indicates the number of chains of each component, ‘l’ denotes the base *λ* ranging from 0 to 0.3, and ‘F’ denotes the model with a flexible elastic network.

**Figure S9:**
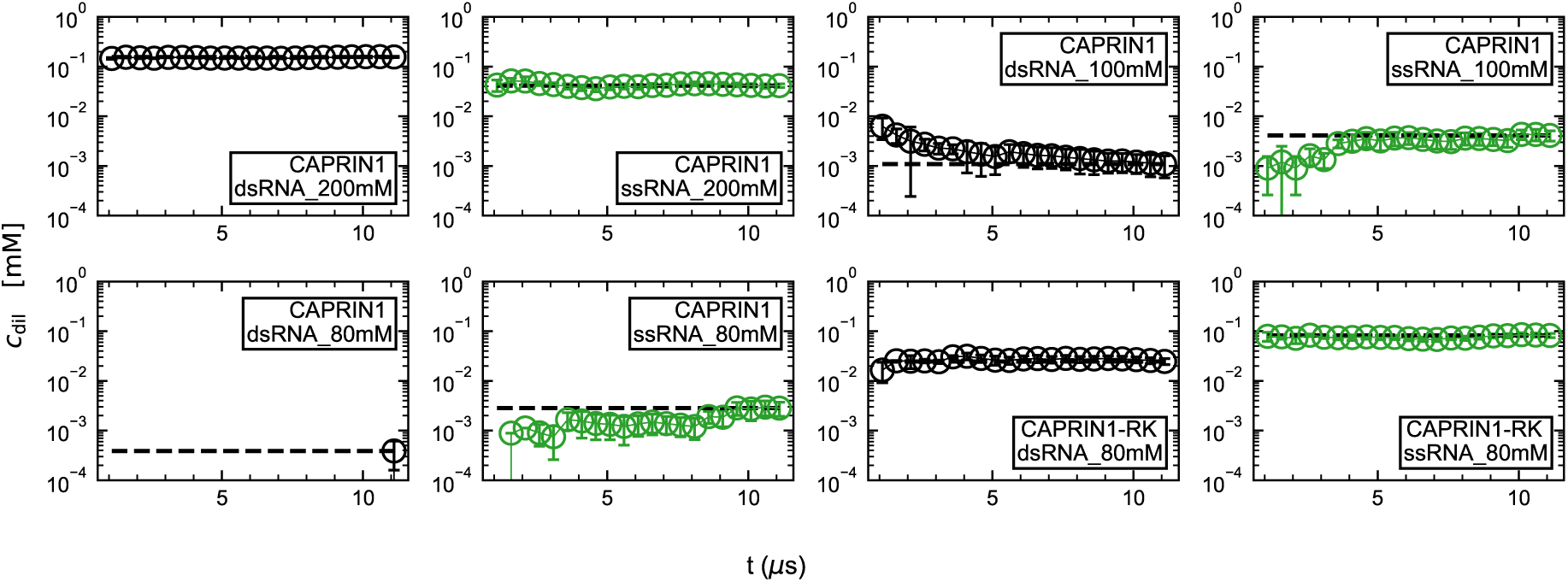
Convergence assessment of the CAPRIN1 systems based on the dilute-phase concentration of the clients (dsRNA or ssRNA). Error bars show the standard error of the mean. CAPRIN1 types (wild type or variant), client types and ionic concentrations are indicated.

**Figure S10:**
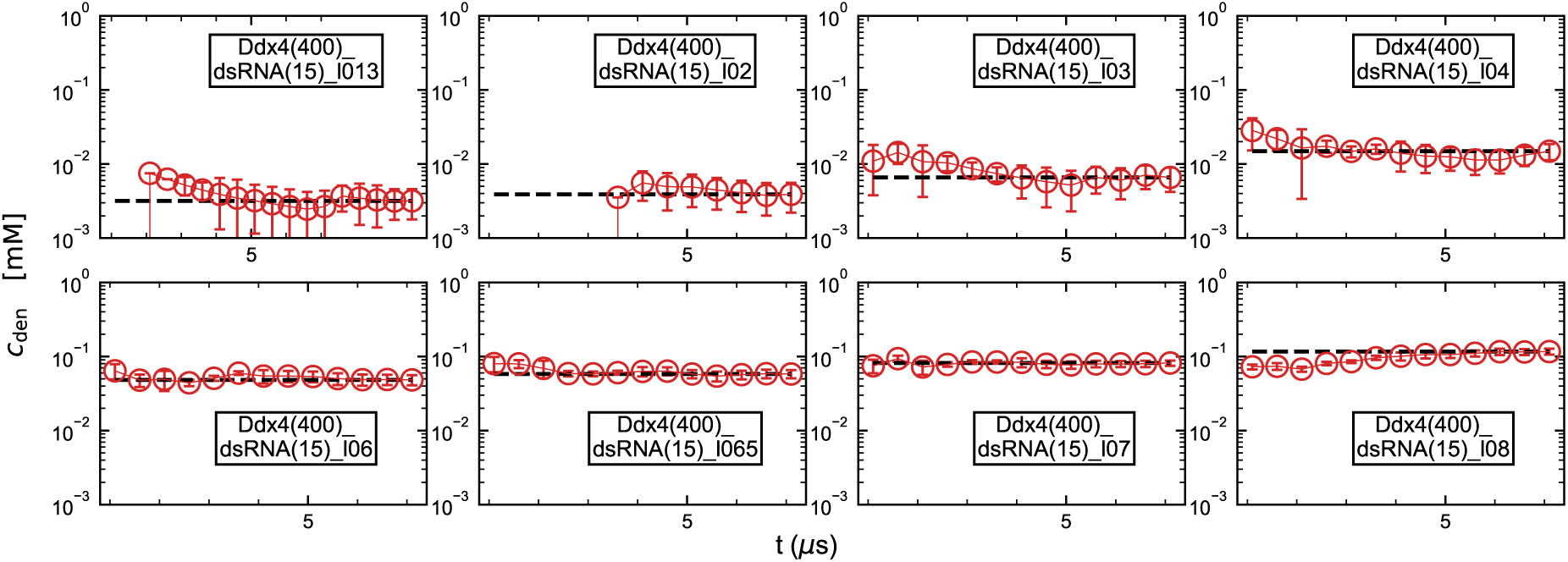
Convergence assessment of the Ddx4N1–dsDNA systems based on the dense-phase concentration of dsDNA. Error bars show the standard error of the mean. The number in parentheses indicates the number of chains of each component, ‘l’ denotes the base *λ* ranging from 0.13 to 0.8.

